# The development and validation of long-read ITS-1/5.8S/ITS-2 nemabiome metabarcoding for ovine gastrointestinal nematodes using Oxford Nanopore Technologies (ONT) sequencing

**DOI:** 10.64898/2025.12.04.692384

**Authors:** Eléonore Charrier, Rebecca Chen, Elizabeth Redman, Sawsan Ammar, Camila Meira, Camila Queiroz, John. S. Gilleard

## Abstract

ITS-2 rRNA nemabiome metabarcoding is increasingly used to characterize gastrointestinal nematode (GIN) communities. While powerful, current approaches have some limitations in their flexibility and applicability to smaller-scale studies and diagnostic use. The short read lengths provided by the Illumina platform may lack discriminatory power for some closely related species and also pose a challenge for new marker selection and primer design. To address these challenges, we have developed ITS-1/5.8S/ITS-2 rRNA Oxford Nanopore Technologies (ONT) long-read metabarcoding for ovine gastrointestinal nematodes. Samples from two previous field studies, from UK and western Canadian sheep farms were used. ITS-1/5.8S/ITS-2 long-read metabarcoding showed strong concordance with prior ITS-2 data for the major GIN species in both datasets, with minor discrepancies for some low abundance taxa mainly due to differences in reference sequence database representation.

We also used the PrimerTC tool to design a new primer pair, EC1 and EC2, to minimize the amplification of off-target fungal sequences derived from fecal DNA and maximize the nematode sequence read depth when ONT ITS-1/5.8S/ITS-2 metabarcoding was applied directly to ovine fecal stool DNA.

In summary, ITS-1/5.8S/ITS-2 ONT long-read nemabiome metabarcoding showed good agreement with ITS-2 metabarcoding and the use of primer pair EC1/EC2 should make the approach more tolerant of fecal contamination of parasite material, with the potential for direct application to ovine fecal DNA. Overall, these new developments should make nemabiome metabarcoding more accessible, discriminating, and flexible for both research and diagnostic applications.

## Introduction

Grazing livestock harbor complex communities of co-infecting gastrointestinal nematode (GIN) species (Zajac and Garza 2020; Burgess et al. 2012). Among these, the suborder Strongylida within Clade V of the nematode phylum contains the most prevalent and important group of ruminant gastrointestinal nematodes (Mederos et al. 2010; Blaxter and Koutsovoulos 2015; O’Connor, Walkden-Brown, and Kahn 2006; Bresciani et al. 2017). Due to the variability of pathogenicity, clinical signs, inherent drug sensitivity and acquired drug resistance status (Besier et al. 2016; Mekonnen 2021; Tariq et al. 2008), it is important to know the relative abundance of each nematode species in fecal samples.

Traditional approaches for monitoring gastrointestinal nematodes in ruminants involve time-consuming techniques that are not ideal for quick, routine diagnostics. Most commonly, fecal egg counts are employed to assess GIN levels in livestock which is traditionally done using microscopy-based methods whereby parasite eggs are counted in fecal samples (Sabatini et al. 2023; Gordon and Whitlock 1939; Vercruysse and Claerebout 2001). However, direct microscopic visualization of eggs cannot distinguish between the different gastrointestinal strongyle species affecting ruminants (Sabatini et al. 2023) and further culturing and subsequent morphological examination of third-stage larvae (L3) is needed to achieve this goal (Knoll et al. 2021; Roeber, Jex, and Gasser 2013c; Gasser et al. 2008). Consequently, it is necessary to either identify these parasites through morphological examination of third-stage larvae following coproculture or use molecular techniques for identification on eggs or hatched larvae (Gasser et al. 2008; Roeber, Jex, and Gasser 2013a; Knoll et al. 2021). However, the conditions employed for nematode egg development during coproculture may introduce a potential bias in the species composition of the resulting L3 populations (Roeber and Kahn 2014). Additional challenges associated with microscopic diagnostic methods are that CERBM-GIE IGBMC they are time-consuming and labour-intensive (Gasser et al. 2008; Roeber, Jex, and Gasser 2013a; Roeber and Kahn 2014; Nielsen 2021; Borkowski et al. 2020) making them difficult to apply to a large number of samples or for rapid diagnostic purposes. Furthermore, the identification of L3s by morphology requires specialist expertise which is often unavailable. The accuracy obtained using molecular techniques, such as real-time PCR (Roeber and Kahn 2014; Learmount et al. 2009) and ddPCR (Elmahalawy et al. 2018) approaches, is higher, and more suitable to large-scale applications but achieving precise quantitation requires thorough optimization and validation in each laboratory (Roeber, Jex, and Gasser 2013a;). Additionally, individual validation of each species-specific assay is required, limiting the identification to only those species expected to be present and for which such validation has been performed.

ITS-2 nemabiome metabarcoding, using Illumina short-read sequencing, was developed to provide an alternative approach to the identification and relative quantitation of gastrointestinal nematode species in fecal samples (Avramenko et al. 2015; Queiroz et al. 2020; Redman et al. 2019). This method is increasingly used in a variety of host species such as cattle, sheep, cervids, bison and horses and the ITS-2 rRNA marker provides good resolution for most of the most important strongylid gastrointestinal nematode species found in these hosts (Gasser et al. 2008; Avramenko et al. 2015; 2017; 2018; Poissant et al, 2021; Queiroz et al. 2020; Zarlenga et al. 2001; Gasser 2006). However, this method still has some limitations. ITS2 lacks resolution for certain closely strongylid GIN species and is not necessarily the best genetic marker for nematodes from other phylogenetic groups of nematodes (Ramünke et al. 2018 and Diekmann et al. 2025). Furthermore, the short sequence read lengths provided by the Illumina platform restrict flexibility to target other genetic markers both in terms of primer design and the ability to maximize the number of potential variable sites. The Illumina platform is also not always locally available and there is often a need to batch large numbers of samples to make the approach cost effective which further reduces its flexibility for smaller project and laboratory use. Hence, ITS2 nemabiome short-read metabarcoding does not provide a comprehensive solution to metabarcoding of all nematode groups in all hosts for all projects.

Different parts of the rRNA cistron vary in their evolutionary divergence and have different, only partially overlapping, representations in reference sequence databases (Charrier et al, 2024; Hillis, David, Dixon 1991). Consequently, the ability to target different regions of the rRNA cistron would broaden the species coverage and also potentially improve the accuracy of nemabiome metabarcoding. However, these targets are too long for paired-end Illumina short-read sequencing. Consequently, here we have explored the use of Oxford Nanopore Technologies (ONT) sequencing for nemabiome metabarcoding to allow a wider range of genetic markers to be targeted. The ONT platforms such as MinION, Flongle, GridION and PromethION also provide more flexible systems suitable not just for larger research and surveillance projects but also for routine diagnostic applications on smaller sample sets.

Another limitation of the existing nemabiome metabarcoding methods is that they involve preparing DNA from parasites that have been purified from fecal samples, including eggs, hatched L1s, or coprocultured L3s. Eliminating the requirement to purify parasites from fecal samples and enabling the direct application of nemabiome metabarcoding to stool DNA would offer a significant advantage in terms of speed, thereby enhancing its use in routine diagnostic protocols. In addition to some general challenges to PCR amplification of parasite templates directly from fecal DNA such as the physical resilience of eggs and the presence of PCR inhibitors, a particular challenge for metabarcoding is the number of off-target sequences amplified which can seriously compromise read depth (Harmon et al. 2007; Ayana et al. 2019; Roeber, Jex, and Gasser 2013b). This can also be a problem for harvested parasite stages still contaminated with fecal debris.

In this current study we have developed ONT sequencing-based nemabiome metabarcoding for ovine gastrointestinal parasitic nematode communities targeting the ITS-1/5.8S/ITS-2 rRNA marker. We used previously characterized field samples from sheep farms from Western Canada and the United Kingdom to validate ITS-1/5.8S/ITS-2 rRNA ONT nemabiome metabarcoding and found an overall high level of agreement with ITS-2 rRNA Illumina short-read nemabiome metabarcoding data. We have also developed a new primer pair — EC1 and EC2 — designed to minimize off-target amplification when ITS-1/5.8S/ITS-2 rRNA ONT metabarcoding is directly applied to ovine fecal (stool) DNA or fecally contaminated parasite samples. These developments should improve the flexibility of nemabiome metabarcoding and make it more suitable for smaller scale research projects and diagnostic applications.

## Methods

### 2.1 Re-analysis of ITS-2 rRNA Nemabiome metabarcoding data

The ITS-2 rRNA Illumina short-read metabarcoding data that is used in this study was previously generated (Queiroz et al. 2020; Redman et al. 2019). It was reanalyzed here to compare ITS-2 rRNA and ITS-1/5.8S/ITS-2 rRNA ONT metabarcoding data.

#### 2.1.1 Origin of ITS-2 rRNA Illumina short-read metabarcoding data

Briefly, ovine fecal samples were collected from ewes from 90 flocks across Western Canada between 2014 and 2018, and 119 farms within the United Kingdom in 2008 (Queiroz et al. 2020; Redman et al. 2019). First stage larvae were harvested from pools of fecal samples comprising of 20 ewes per farm. Genomic DNA lysates were made for each farm pool from between 200 and 1000 L1s for Western Canadian and United Kingdom samples respectively and 1:10 or 1:40 dilutions were used as templates for ITS-2 rRNA nemabiome metabarcoding using the Illumina MiSeq platform (Queiroz et al. 2020; Redman et al. 2019).

#### 2.1.2 Bioinformatic and Statistical re-analysis of ITS-2 rRNA Illumina short-read metabarcoding data

The original fastq files from several different Illumina ITS-2 metabarcoding assays were re-analyzed using a bioinformatic pipeline (Beaumelle et al. 2021) based on DADA2 software version 1.20.0 (Callahan et al. 2016) on R studio version 4.1.1. As per DADA2 developer recommendation (https://github.com/benjjneb/dada2/issues/1652), the reads from each run were processed separately, to reduce error rates. Using the default parameters of Cutadapt version 3.5 (Martin 2011) the primers were detected and removed from classified sequences. Additionally, the following reads were removed; those shorter than 50 bp, those with a maximum expected error >1 for the forward reads or >2 for the reverse sequences, or those with a quality score of two or lower. The filtered and merged reads from each separate run were then combined for the subsequent bioinformatic analysis and the DADA2 software was used to remove chimeric sequences. The remaining sequences were then classified taxonomically using the IDTaxa classification method with a confidence threshold set at 60 against a nematode ITS-2 rRNA reference sequence database available at www.nemabiome.ca (Workentine et al. 2019). Additionally, samples with a total mapped read depth of lower than 1000 reads were removed from downstream analysis. The relative abundance of each species was calculated by dividing the total number of reads for each species by the total number of reads from each sample. Correction factors were not applied in this study as the aim was to directly compare the outputs of Illumina and ONT nemabiome metabarcoding. Results of the taxonomic classification and relative quantitation of nematode species were visualized in R Studio using the ggplot2 package.

### 2.2 ONT rRNA ITS-1/5.8S/ITS-2 long-read metabarcoding using the MinION device

#### 2.2.1 ONT long-read metabarcoding of the ITS-1/5.8S/ITS-2 rRNA locus using the NC5/NC2 primer pair

The same genomic DNA samples from UK and Western Canadian ovine GIN populations, that were originally used as templates for ITS-2 Illumina metabarcoding, were used for ONT metabarcoding (Queiroz et al. 2020; Redman et al. 2019). We used 8µL of 1/10 dilutions of template DNA in the first round of PCR amplifications of the ITS-1/5.8S/ITS-2 marker. NC5-Np-Adp (forward) and NC2-Np-Adp (reverse) primers were synthesized comprising the previously described NC5 and NC2 primers which flank the ITS-1/5.8S/ITS-2 region (Gasser et al. 1993; Newton et al. 1998) but with ONT sequencing adapter tags (https://community.nanoporetech.com/docs/prepare/library_prep_protocols/pcr-96-barcoding-amplicons/v/pbac96_9069_v109_revv_14aug2019/pcr-barcoding-amplicons-cdna?devices=minion) on their 5 ’end (Supplementary Table 1). NC5 and NC2 are complementary to the 3 ’terminus of the 18S gene and 5 ’terminus of the 28S gene coding sequences respectively and amplify a 700-1000 bp product from the major ovine strongylid GIN species. The NC5-Np-Adp and NC2-Np-Adp primers were used to amplify the ITS-1/5.8S/ITS-2 region of the rRNA cistron for Nanopore long-read sequencing. PCR conditions were 10 µL of KAPA HiFi Fidelity Buffer (5x), 1.5 µL of dNTPs (10mM), 1.5 µL NC5 Primer (10 µM), 1.5 µL NC2 Primer (10 µM), 1 µL KAPA HiFi HotStart DNA Polymerase, 30.5 µL ddH2O and 4 µL diluted lysate. The thermocycling parameters were 95°C for three minutes, 35 cycles of 98°C for 20 seconds, 58°C for 15 seconds, and 72°C for 30 seconds followed by a final extension of 72°C for two minutes. PCR products were purified with AMPure XP Magnetic Beads (1X) (Beckman Coulter, Inc). The purified PCR products were multiplexed using the Oxford Nanopore barcoded primers (Oxford Nanopore, EXP-PBC096). The following PCR conditions were used: 1 µL Barcoded Primer (one BC01 – BC96 at 10 µM), 25 µL LongAmp Hot Start Taq Master Mix (2X) and 24 µL of purified first-round PCR product as template. The thermocycling parameters were 95°C for three minutes, 13 cycles of 95°C for 15 seconds, 62°C for 15 seconds, and 65°C for 50 seconds followed by a final extension of 65°C for five minutes. The PCR products were purified using AMPure XP Magnetic Beads (1X). The concentration of each PCR sample product was assessed using Qubit Fluorometric Quantification (ThermoFisher Scientific, Inc). A master sequencing library was created by pooling 1.5 µg of each purified product. The NEBNext Ultra II End Repair / dA-tailing Module (New England BioLabs, E7546) was then used for the End-prep of the pooled library, followed by the ligation of ONT-specific adapters (Oxford nanopore, SQL-LSK112). The final concentration of the pooled library was assessed by Qubit fluorometric quantification. The prepared pooled library was run on an ONT Mk1B sequencer at a concentration of 50 fmol. The MinION device was set to generate only fast5 files with no post-run analysis. All protocols were carried out using Oxford Nanopore’s standard operating protocol (https://community.nanoporetech.com/docs/prepare/library_prep_protocols/ligation-sequencing-amplicons-sqk-lsk114/v/acde_9163_v114_revq_29jun2022).

#### 2.2.2 Bioinformatic and statistical analysis of ITS-1/5.8S/ITS-2 rRNA ONT metabarcoding data

We developed a custom bioinformatic pipeline to analyze the ONT metabarcoding data (Figure 1). Reads from each sample were basecalled (using the super-accuracy parameter) and de-multiplexed, post-sequencing run, using the Oxford Nanopore Technologies basecaller software Guppy version 6.1.7. Additionally, the barcode sequences were trimmed during this basecalling and de-multiplexing step. First, primer sequences were identified and removed using Cutadapt software version 4.1 (Martin 2011). Reads from each sample were then filtered using Filtlong software version 0.2.1 (https://github.com/rrwick/Filtlong) with the following parameters: 1) reads shorter than 700 bp were removed; 2) the lowest 10% of reads based on quality score were removed; 3) reads with a mean quality weight of <20 were removed (chosen based on the Oxford Nanopore Q20 chemistry kit used for sequencing). Finally, the remaining reads for each sample were classified taxonomically and assigned against a database of ITS-1/5.8S/ITS-2 reference sequences (Charrier et al, 2024) using the Minimap2 software version 2.23 (Li 2018). Samples with a total read number of less than 1000 were removed. The relative abundance of each species was calculated by dividing the total number of reads for each species by the total number of reads from each sample.

**Figure 1.**
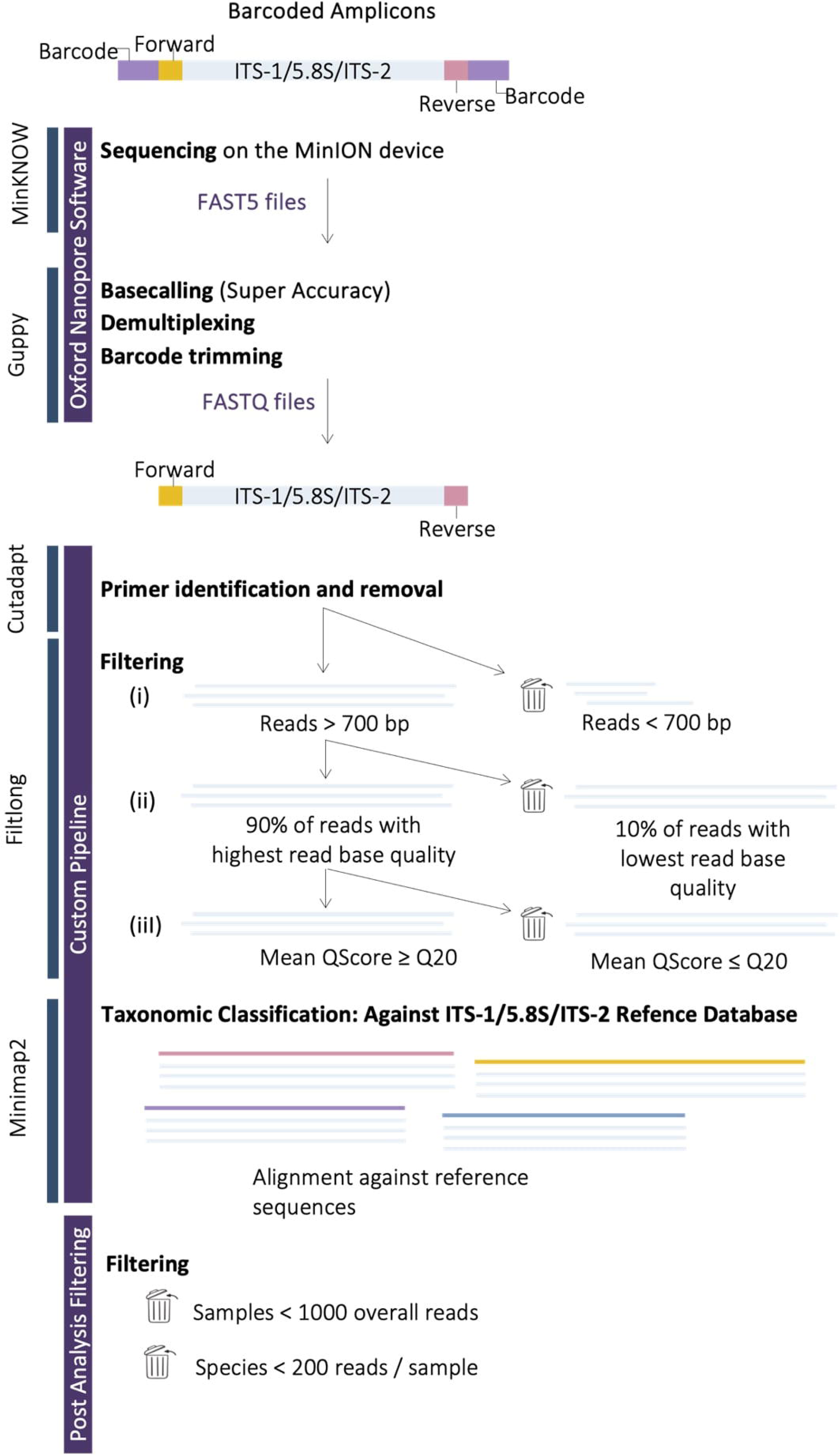
Flowchart of the ITS-1/5.8S/ITS-2 ONT long-read nemabiome metabarcoding bioinformatic analysis pipeline.

Lin’s Concordance Correlation Coefficient was calculated using the CCC {DescTools} package in R studio. CCC values > 0.8 are classified as a near-perfect agreement (CCC = 1, which would represent a perfect agreement). Data were analyzed using a Kruskal-Wallis test with geographical region as a factor. The analysis was done in R using the *kruskal.test()* function (https://www.rdocumentation.org/packages/stats/versions/3.6.2/topics/aov). Statistical significance was set at p < 0.05.

### 2.3. Developing ITS-1/5.8S/ITS-2 rRNA metabarcoding primers that minimize off-target amplification from fecal stool DNA template

#### 2.3.1 Sample Collection and processing of fecal samples

Fecal samples were collected from individual ewes in Western Canada (Saskatchewan, Alberta, and British Columbia) between 2017–2019 and in 2023. Producers collected fresh samples, placed them in plastic bags with the air removed, and shipped them at room temperature. Samples were stored at 15 °C in the lab and processed within 5 days. For each flock, 15–20 samples were pooled (10–32 g per sample) and mixed manually. Fecal egg counts (FEC) were determined as the mean of three 2 g aliquots per pool using a modified McMaster method with a three-chamber slide, with a sensitivity of 16.66 eggs per gram (EPG) (Paracount-EPG™, Chalex, LCC– https://www.vetslides.com/product-page/paracount-epg-fecal-analysis-kit-3-chamber-slide). Multiple 0.5 g aliquots of each flock pool were mixed with 1 mL of 95% ethanol and archived at - 80 °C for future DNA preparation.

#### 2.3.2 Preparation of genomic DNA from fecal stool samples

Genomic DNA was isolated from the archived ethanol-fixed fecal samples (0.5g of feces mixed with 1mL of 95% ethanol) using one of the following two DNA isolation methods (i) QIAmp Power Fecal Pro DNA Kit Ethanol-fixed stool samples (500 µL) were washed three times by centrifugation with molecular-grade water, resuspended in 250 µL water, and DNA extracted using the QIAmp Power Fecal Pro DNA Kit (Qiagen, Inc.) following the manufacturer’s protocol. DNA was eluted in 100 µL elution buffer and stored at -20 °C. (ii) Extended freeze/thaw protocol before the ‘QIAmp Power Fecal Pro DNA Kit’ (Venkatesan 2023). Samples (500 µL) were washed as above, then subjected to three freeze-thaw cycles: snap freezing in liquid nitrogen, heating at 100 °C for 15 min, and bead beating for 3 min. DNA was extracted with the QIAmp Power Fecal Pro DNA Kit, eluted in 100 µL, and stored at -20 °C.

#### 2.3.3 Designing ITS-1/5.8S/ITS-2 rRNA PCR primers to minimize off-target amplification from fecal gDNA

Primer Tc was used to design primers EC1 and EC2 (sequences in Supplementary Table 1) to amplify the nematode ITS-1–5.8S–ITS-2 rDNA locus whilst minimizing off-target amplification of fungal sequences from fecal DNA (Charrier et al., 2024). Primer design involved two steps: (1) assessing sequence conservation against nematode ITS1/5.8S/ITS2 rDNA sequences to ensure broad taxonomic coverage across the phylum, and (2) checking against fungal rDNA sequences (NCBI fungal 18S and 28S databases) to minimize amplification of fungal DNA. EC1, a forward primer targeting the 3’ end of 18S rDNA, shows high identity across 514 nematode genera from 2,645 reference sequences representing 1,391 species. EC2, a reverse primer targeting the 5’ end of 28S rDNA, shows high identity across 117 genera from 254 reference sequences representing 204 species. Both primers show minimal identity to 321 fungal 18S and 113 fungal 28S sequences (https://ftp.ncbi.nlm.nih.gov/refseq/TargetedLoci/Fungi/) (Table 1).

**Table 1.**
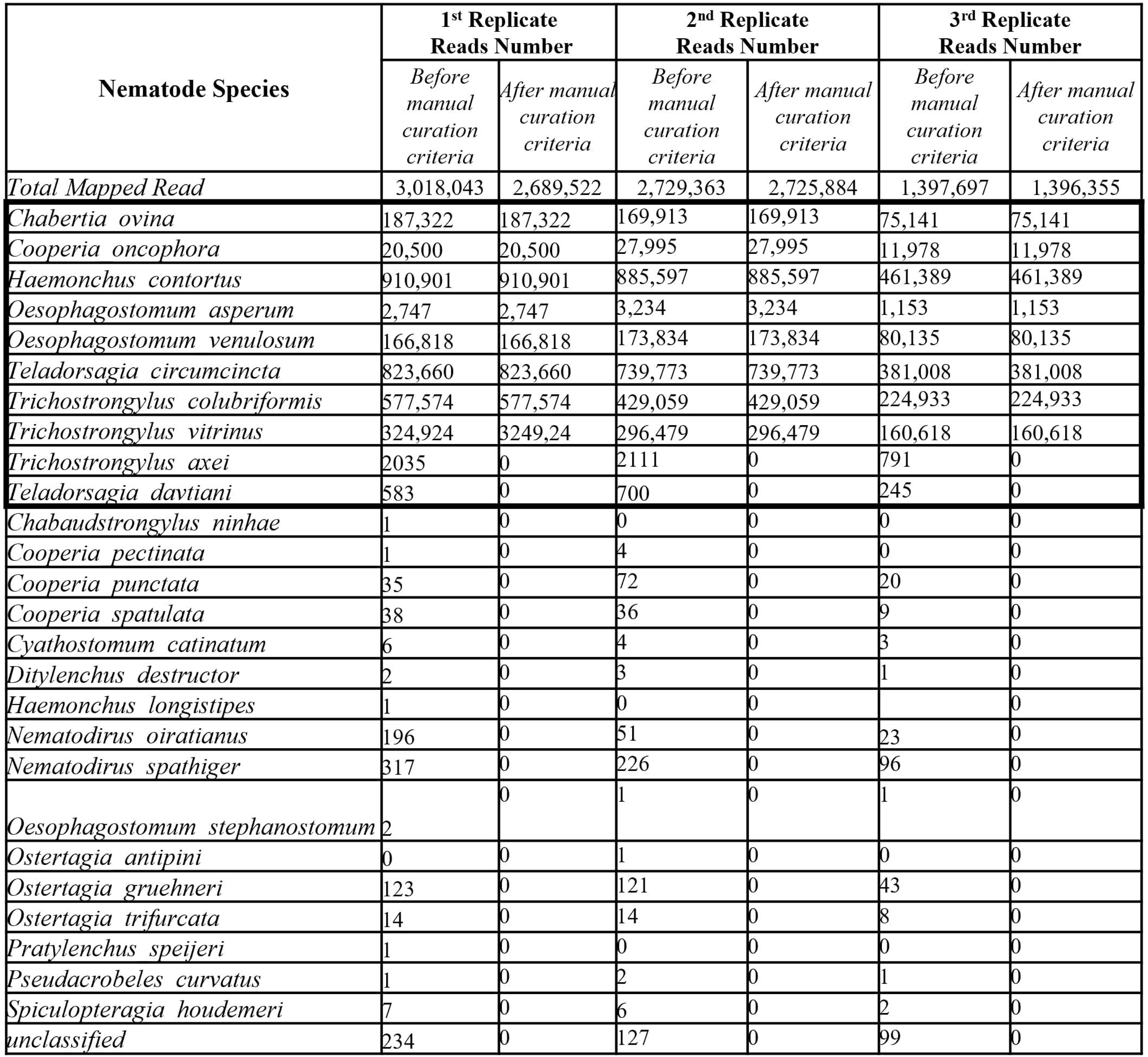
The numbers of ONT ITS-1/5.8S/ITS-2 rDNA sequence reads mapping to each nematode species before and after applying manual curation filtering criteria. The ITS-1/5.8S/ITS-2 rDNA was independently amplified from each GIN L1 pool three separate times to produce three independent replicate libraries each sequenced on the ONT MinION for 30 hours. Before the application of manual curation criteria, species and sequencing reads were obtained through the ITS-1/5.8S/ITS-2 nemabiome metabarcoding bioinformatic pipeline. After application of manual curation criteria, results were obtained after species with less than 200 overall mapped reads and samples with less than 1000 mapped reads were removed.

#### 2.3.4 ONT ITS-1/5.8S/ITS-2 rRNA metabarcoding from ovine fecal DNA using the EC1/EC2 primer pair

EC1-Np-Adp (forward) and EC2-Np-Adp (reverse) primers were synthesized by adding ONT sequencing adapter tags to the 5’ ends of the previously developed EC1 and EC2 primers (https://community.nanoporetech.com/docs/prepare/library_prep_protocols/pcr-96-barcoding-amplicons/v/pbac96_9069_v109_revv_14aug2019/pcr-barcoding-amplicons-cdna?devices=minion) (Supplementary Table 1). These primers were used to amplify the ITS-1/5.8S/ITS-2 rRNA region for ONT MinION long-read sequencing. PCR reactions contained 10 µL KAPAHiFi Fidelity Buffer (5x), 1.5 µL dNTPs (10 mM), 1.5 µL of each EC1 and EC2 primers (10 µM), 1 µL KAPA HiFi HotStart DNA Polymerase, 0.2 µL BSA (20 mg/µL), 29.3 µL ddH₂O, and 5 µL lysate. Thermocycling was 95 °C for 3 min; 35 cycles of 98 °C for 20 s, 58 °C for 15 s, 72 °C for 1 min; and a final extension at 72 °C for 2 min. Subsequent Nanopore ITS-1/5.8S/ITS-2 metabarcoding and analysis followed previously described protocols.

## Results

### 3.1. Determination of relative species abundance in ovine gastrointestinal communities by ONT ITS-1/5.8S/ITS-2 rRNA long-read nemabiome metabarcoding

#### 3.1.1 Assessment of intra- and inter-species variation of ITS-1/5.8S/ITS-2 rRNA reference sequences

We have previously developed a non-redundant database of full-length nematode ITS-1/5.8S/ITS-2 rRNA reference sequences specifically to support ONT long-read nemabiome metabarcoding available at www.nemabiome.ca (Charrier et al, 2024). Analyzing the 10,107 sequences within this new database revealed that the ITS-1/5.8S/ITS-2 rRNA marker ranges in size as follows; 702 - 1,700bp across the Phylum Nematoda, 706-1,500bp across the order Strongylida and 706-1,205bp across the superfamily Trichostrongyloidea (Supplementary Table 2).

For the most important domestic ruminant parasite species of interest within the superfamily *Trichostrongyloidea - Chabertia ovina, Cooperia pectinata, Cooperia oncophora, Cooperia punctata, Ostertagia ostertagi, Ostertagia trifurcata, Ostertagia venulosum, Oesophogostumum asperum, Oesophagostomum columbianum, Trichostrongylus probolurus, Trichostrongylus axei, Trichostrongylus colubriformis, Trichostrongylus vitrinus, Teladorsagia davtiani, Teladorsagia circumcincta* - we used all available sequences in the ITS-1/5.8S/ITS-2 database to assess both the intra- and inter-species variation. The results suggest that the marker provides good discrimination for all these species. *Cooperia oncophora* and *Cooperia punctata*, the most closely related species, exhibit an inter-species identity ranging from 96.5% to 98%. There is a range of 55 to 110 nucleotide difference between these two species (Figure 2B). Similarly, the inter-species identity between *Trichostrongylus axei* and *Trichostrongylus colubriformis*, two other closely related species, falls within the range of 87.9% to 97.4%, (Figure 2A), with a nucleotide difference between 20 and 83 (Figure 2B).

**Figure 2.**
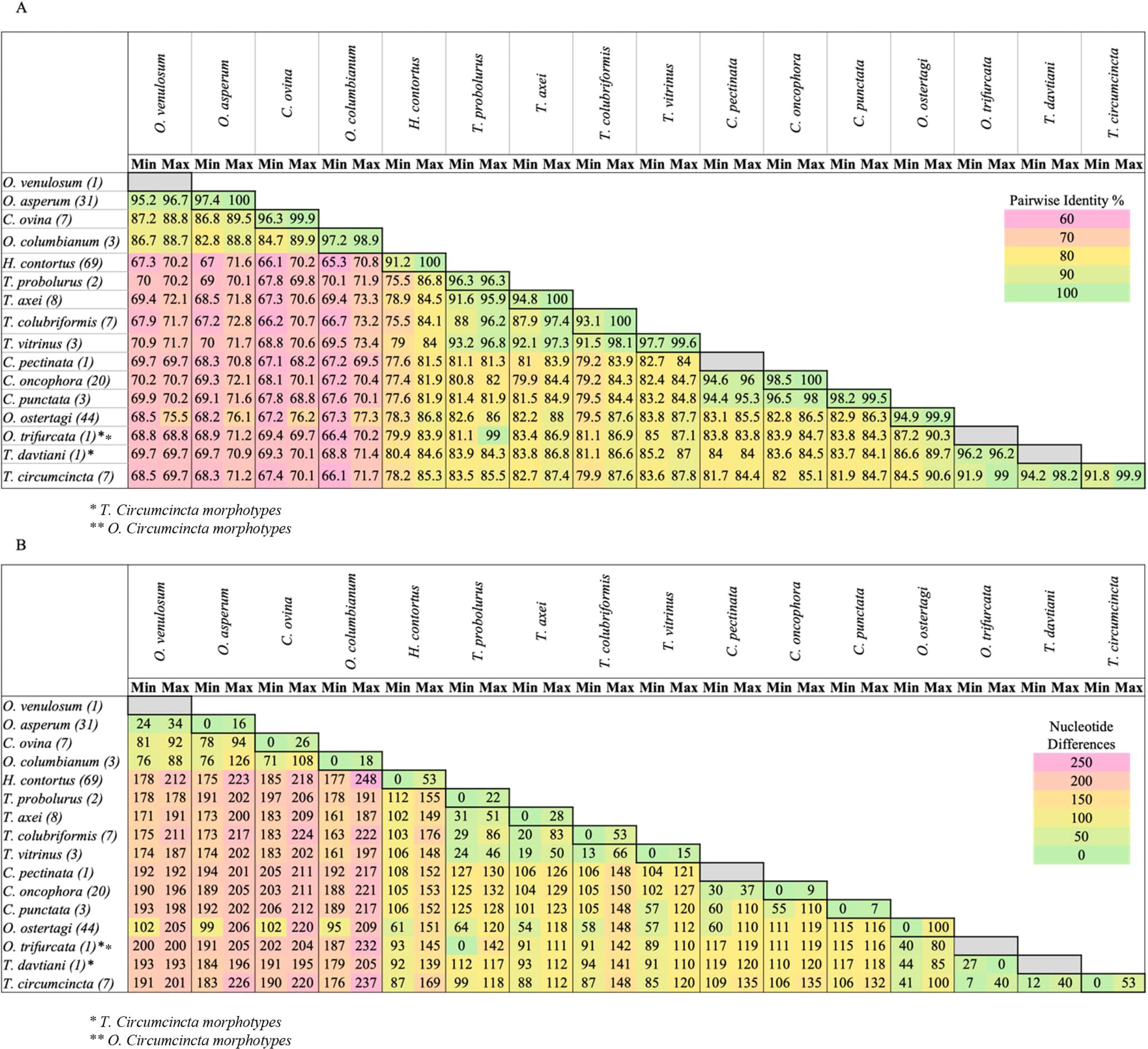
Inter- and intra-species variation of ITS-1/5.8S/ITS-2 sequence identity. All full-length ITS-1/5.8S/ITS-2 sequences, currently available in the ITS-1/5.8S/ITS-2 database (nemabiome.ca: https://staging--nemabiome.netlify.app/), were compiled for each species. (A) The minimum and maximum percent sequence within and between each species is indicated in each cell. The number of unique full-length sequences for each species in the database is indicated in parenthesis next to the species name. Sequences were aligned with MUSCLE alignment (Edgar 2004) using the default parameters using Geneious version 10.1.3. (B) The lowest and highest number of nucleotide differences between sequences for each species. The number of unique full-length sequences for each species in the database is indicated in parenthesis next to the species name. Sequences were aligned with MUSCLE (Edgar 2004) using the default parameters using Geneious version 10.1.3. An absolute pairwise character difference matrix was generated using the *haplotypes::distance()* R package.

#### 3.1.2 Assessment of ITS-1/5.8S/ITS-2 rRNA long-read ONT nemabiome metabarcoding

We tested ITS-1/5.8S/ITS-2 rRNA long-read ONT nemabiome metabarcoding on sets of ovine gastrointestinal parasitic nematode populations previously collected from 34 Western Canadian and 43 UK sheep flocks. These populations consisted of pools of L1 larvae derived from eggs harvested from fecal matter by flotation (pooled from 20 ewes per flock) and so were largely free of contaminating bacteria, fungi, or other material. Here, we applied ITS-1/5.8S/ITS-2 rRNA long-read ONT metabarcoding to the same genomic DNA template preparations that were previously used for ITS-2 short-read Illumina metabarcoding (Redman et al. 2019; Queiroz et al. 2020).

Primers NC5-Np-Adp and NC2-Np-Adp were used to PCR amplify the ITS-1/5.8S/ITS-2 marker from each genomic DNA preparation three separate times to produce three independent replicate ONT sequencing libraries. These libraries were sequenced on the MinION device for 30 hours to generate 6.48, 5.87 and 3.22 million total reads for each of the three libraries respectively. The raw sequencing data from each replicate library was independently processed through our nemabiome metabarcoding analysis pipeline (Figure 1). These custom pipeline filtering steps resulted in 3.018, 2.729 and 1.397 million reads passing the filtering steps for each of the triplicate libraries respectively.

Out of these reads only 234, 127 and 99 reads, did not map to any reference sequence in the nematode ITS-1/5.8S/ITS-2 database respectively with the remainder mapping to a total of twenty-six nematode species across the three library replicates (Table 1). With the exception of *Cooperia oncophora*, the nematode species with at least 100 sequence reads are known to be ovine parasites. However, there were multiple species with very low read counts that are not expected to parasitize sheep.

We subjected the sequencing reads outputted by the pipeline to additional manual curation by removing any samples with less than a total of 1,000 reads in any of the three replicates and removing any species with less than 200 reads across all samples. This manual curation resulted in final total read numbers of 3.017, 2.729 and 1.397 million reads respectively for each of the three replicate libraries and reduced the list of nematode species with mapped reads from twenty-six to ten (Figure 3 and Table 1). The data revealed some differences in the GIN communities from the two geographical regions (Figure 3). Most notably, *H. contortus* is much more prevalent and abundant in the Canadian than the UK flocks; 34/34 (100%). Canadian flocks were all positive for *H. contortus* with an overall relative abundance of 43.2% compared to 37/43 (86%) positive UK flocks with an overall relative abundance of 21.5% (Figure 3A). Conversely, *T. vitrinus* was much less prevalent and abundant in the Canadian than the UK flocks; all flocks were positive for *T. vitrinus* but with an overall relative abundance of 3.1% in the Canadian Flocks compared to an overall relative abundance of 16.8% for the UK farms. There were also notable differences in the prevalence and relative abundance of *T. circumcincta* and *T. colubriformis* with overall abundance in Western Canada of 10.13% and 16.78% respectively and of 25.92% and 3.08% in the United Kingdom samples respectively (Figure 3B).

**Figure 3.**
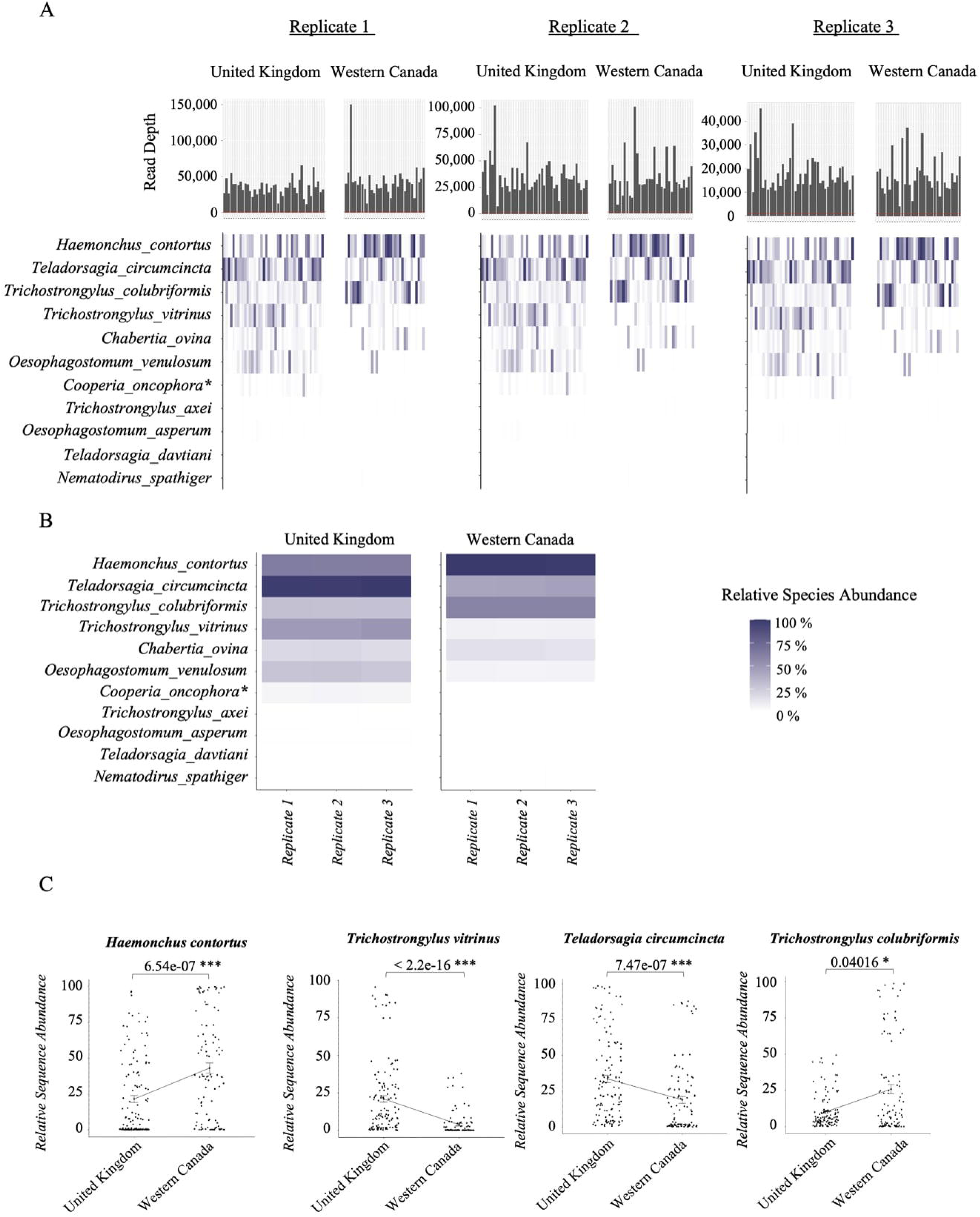
Nematode relative species abundance after post-bioinformatic manual curation of ITS-1/5.8S/ITS-2 rDNA ONT nemabiome metabarcoding data. The ITS-1/5.8S/ITS-2 rDNA was independently amplified from genomic DNA made from each GIN L1 pool three separate times to produce three independent replicate libraries, each of which was sequenced on the Oxford Nanopore MinION for 30 hours. Following analysis by the bioinformatic pipeline (Figure 1), the data was manually curated by the exclusion of samples with a total read depth below 1000 reads and of nematode species with fewer than 200 reads across all samples. The data comprises 43 UK samples and the 34 Western Canadian samples. (A) The top histogram shows the read depth of the mapped nematode ITS-1/5.8S/ITS-2 reads in each sample. The 1000 sequence read threshold is indicated by the red line. The heatmaps show the relative abundance of ITS-1/5.8S/ITS-2 rRNA reads for the different nematode species within each sample for the three replicates after manual curation. (B) The heatmaps show the relative abundance of nematode species within each replicate library after manual curation. *\*C. curticei* is misidentified as *C. oncophora* as the current ITS-1/5.8S/ITS-2 database used does not contain reference sequences for *C. curticei* and identified the reads to the closest related species. (C) Kruskal-Wallis’s test of comparing the relative abundance of *H. contortus* (p-value: 6.54e-07)*, T. vitrinus* (p-value: < 2.2e-16)*, T. circumcincta* (p-value: 7.47e-07), and *T. colubriformis* (p-value: 0.04016) between the 43 UK samples and the 34 Western Canadian samples. For each graph, the mean relative abundance is represented as well as the standard error.

For the four most common GIN species (*H. contortus, T. vitrinus, T. circumcincta* and *T. colubriformis*), an analysis of variance using a Kruskal-Wallis test showed the differences in relative abundance between the Canadian and UK samples was statistically significant (*H. contortus* p-value: 6.54e-07*, T. vitrinus* p-value: <2.2e-16*, T. circumcincta* p-value: 7.47e-07, and *T. colubriformis* p-value: 0.04016). (Figure 3C).

#### 3.1.3 Repeatability of ITS-1/5.8S/ITS-2 long-read Oxford Nanopore nemabiome metabarcoding

There was a high level of repeatability of the ITS-1/5.8S/ITS-2 metabarcoding data across the three independent replicate datasets (Figure 4A). The relative abundance of each species averaged at 0.2% (minimum of 0% and maximum of 0.7%) between the three replicates (mean standard deviations; 0.00% - 0.39%) (Data not shown and a Lin’s Concordance Correlation Coefficient of 0.99 for each of our three comparisons) (Figure 4B).

**Figure 4.**
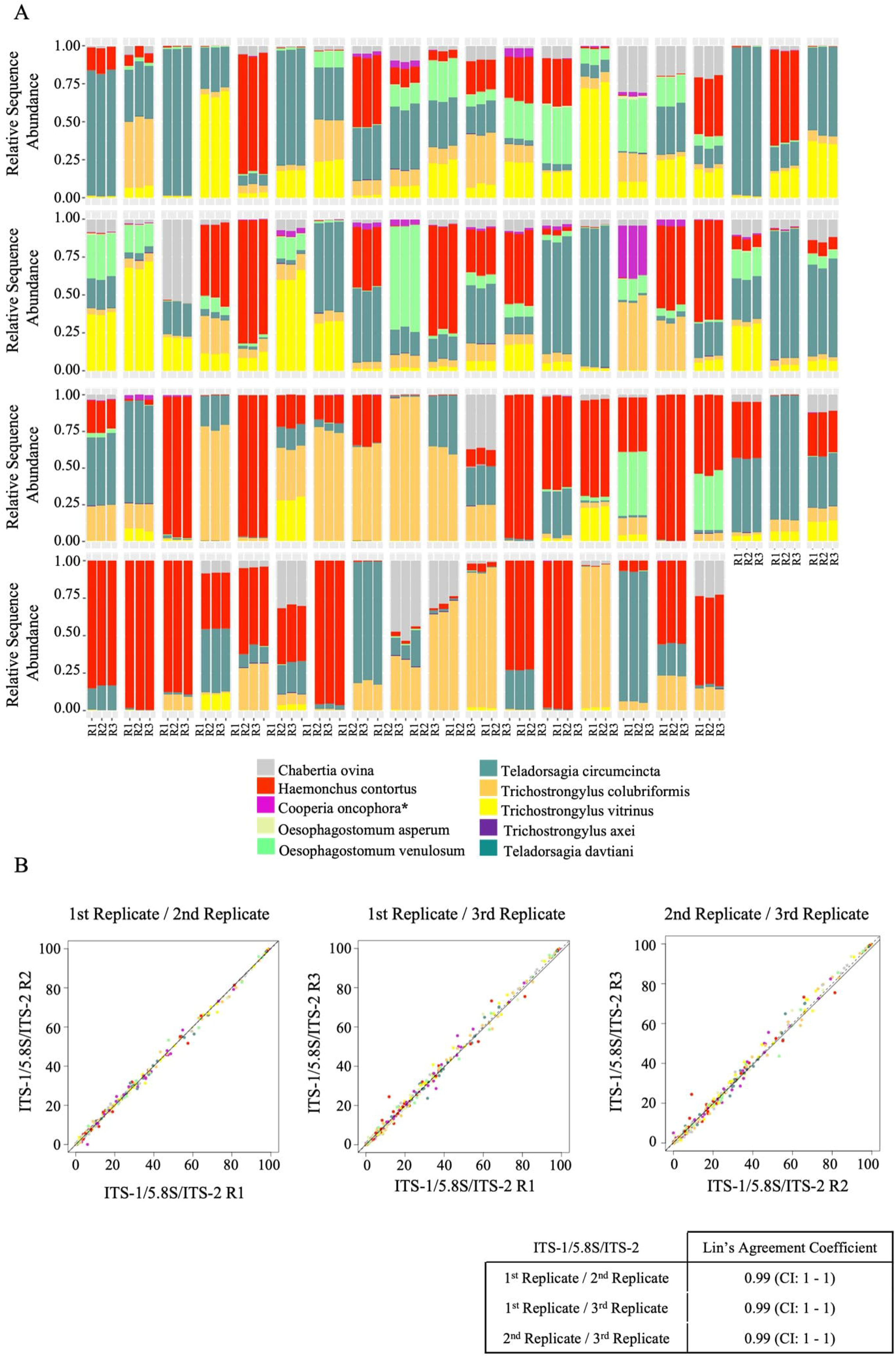
Repeatability of ITS-1/5.8S/ITS-2 long-read ONT nemabiome metabarcoding. The ITS-1/5.8S/ITS-2 rDNA was independently PCR amplified three separate times from each genomic DNA template of pooled L1 larvae to produce three independent replicate libraries sequenced on the ONT MinION device for 30 hours. (A) The chart displays the relative nematode species abundance for each of the three replicates for each sample based on the ITS-1/5.8S/ITS- metabarcoding. (B) The charts plot the abundance of each nematode species in each sample for pairwise comparisons of replicates. The table shows Lin’s Concordance Correlation Coefficient analysis results of the two replicate sequencing libraries being compared. *\*C. curticei* is misidentified as *C. oncophora* as the current ITS-1/5.8S/ITS-2 database used does not contain reference sequences for *C. curticei* and identified the reads to the closest related species.

### 3.2. Comparison of ITS-1/5.8S/ITS-2 rRNA ONT long-read metabarcoding with ITS-2 rRNA Illumina short-read metabarcoding

To ensure a more direct comparison with the ITS-1/5.8S/ITS-2 rRNA Oxford Nanopore metabarcoding, we re-analyzed the ITS-2 rRNA Illumina nemabiome metabarcoding data, previously generated from the same flocks, using an updated DADA2 bioinformatic pipeline (Callahan et al. 2016; Beaumelle et al. 2021). This analysis identified a total of 19 nematode species which reduced to 14 following the same manual filtering criteria applied to the Nanopore metabarcoding data: *C. ovina* (4.07%), *C. curticei* (0.74%), *C. fuelleborni* (0.82%), *C. oncophora* (0.02%), *H. contortus* (26.44%), *O. venulosum* (6.12%), *T. circumcincta* (34.35%), *T. axei* (2.45%), *T. colubriformis* (9.22%), *T. tenuis* (0.03%), *T. vitrinus* (14.41%) as well as *Unclassified Cooperia* (0.13%), *Ostertagia* (0.01%) and *Trichostrongylus* (1.19%) species (Supplementary Table 3).

We compared the ONT long-read ITS-1/5.8S/ITS-2 metabarcoding data with the Illumina short-read ITS-2 metabarcoding data on the same samples. The overall nematode species proportions identified in the three replicate ONT ITS-1/5.8S/ITS-2 rRNA long-read nanopore metabarcoding datasets were very similar to those of the previous short-read Illumina ITS-2 rRNA metabarcoding data (Lin agreement coefficient; 1^st^ replicate = 0.96, 2^nd^ replicate = 0.96, 3^rd^ replicates = 0.96) (Figure 5A and B). There was also minimal discrepancy when considering each of the six major nematode species individual Lin’s agreement coefficient for each of the three replicates; *H. contortus* (0.95, 0.95, 0.95)*, T. colubriformis* (0.94, 0.94, 0.94)*, T. circumcincta* (0.95, 0.96, 0.96)*, T. vitrinus* (0.98, 0.98, 0.98), and *O. venulosum* (0.93, 0.93, 0.93) (Figure 5B and Supplementary Figure 1). However, there was some discrepancy for *Cooperia* species: the Illumina short-read ITS-2 metabarcoding reads mapped to reference sequences of two *Cooperia* species (*C. curticei* and *C. fuelleborni*) whereas the ONT ITS-1/5.8S/ITS-2 long-read metabarcoding reads only mapped to one, *C. oncophora,* resulting in a lower Lin’s agreement (0.81, 0.79, 0.82) (Figure 5B and Supplementary Figure 1). It should be noted there is no *C. curticei* reference sequence in the ITS-1/5.8S/ITS-2 database which is the likely cause of this discrepancy, and this is discussed further below. Also, although both the ONT ITS-1/5.8S/ITS-2 and Illumina ITS-2 metabarcoding identified *T. axei*, the ONT replicates identified this species in a much lower abundance, leading to a very low Lin’s agreement (0.05, 0.06, 0.04) (Figure 5B and Supplementary Figure 1).This discrepancy is also likely due to differences in database representation of this species as is discussed further below.

**Figure 5.**
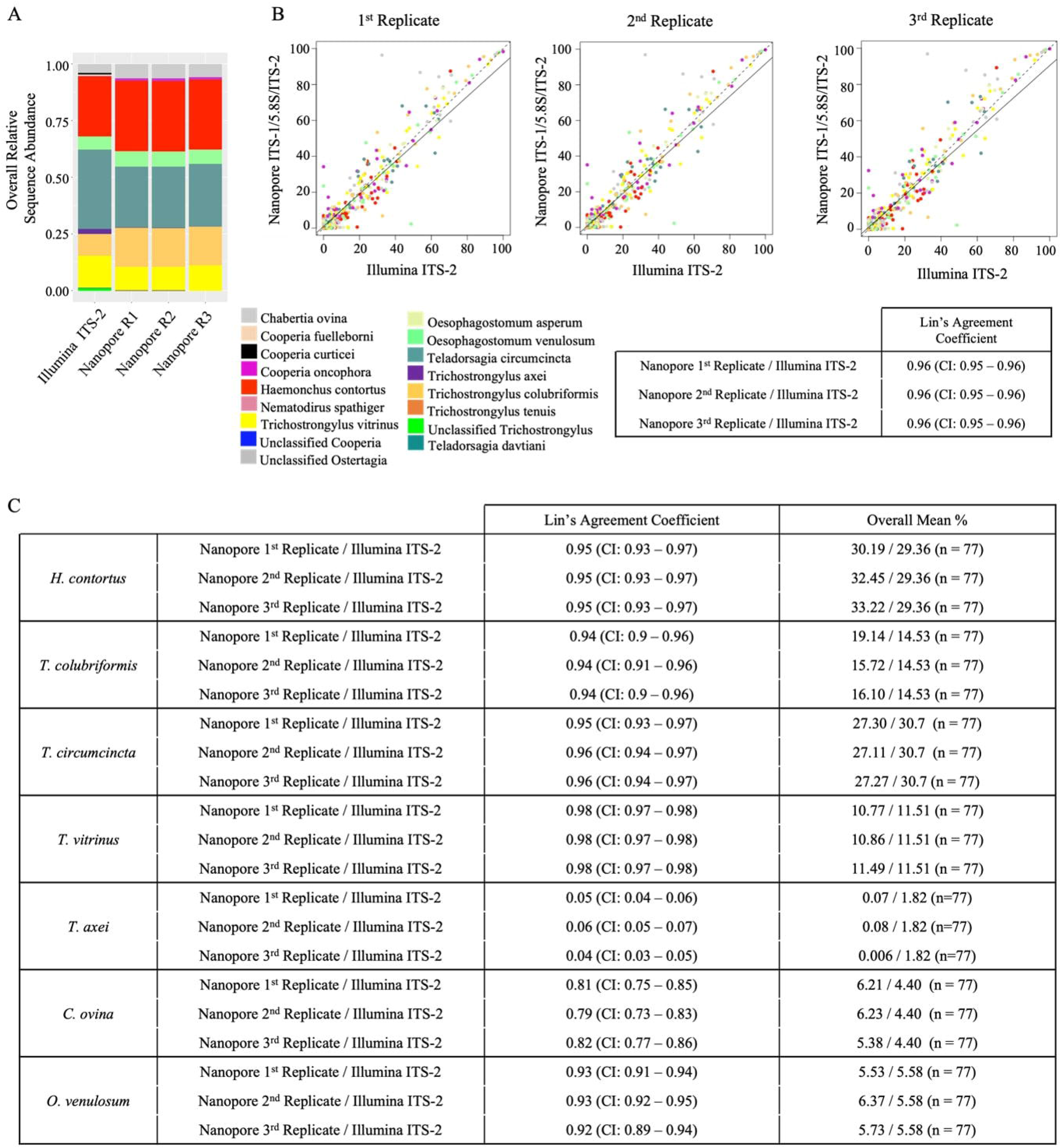
Comparison of ITS-1/5.8S/ITS-2 long-read ONT nemabiome metabarcoding with ITS-2 short-read Illumina metabarcoding data. The ITS-1/5.8S/ITS-2 rDNA was independently PCR amplified three separate times from each genomic DNA template of pooled L1 larvae to produce three independent replicate libraries sequenced on the ONT MinION device for 30 hours. Following analysis and manual curation, this data was compared with ITS-2 rRNA previously generated from the same DNA templates. (A) The chart displays the overall proportional representation of each species for all the ITS-1/5.8S/ITS- metabarcoding and the previously obtained ITS-2 metabarcoding data from all the samples combined. (B) The charts plot the abundance of each nematode species in each sample for pairwise comparisons of each of the ONT ITS-1/5.8S/ITS-2 metabarcoding replicates and the Illumina ITS-2 metabarcoding data. The table shows Lin’s Concordance Correlation Coefficient analysis results of each of the two sequencing libraries being compared. (C) The table shows Lin’s Concordance Correlation Coefficient analysis results of each of the two sequencing libraries being compared for *H. contortus, T. colubriformis, T. circumcincta, T. vitrinus, T. axei, C. ovina* and *O. venulosum*.

### 3.3. Minimizing off-target reads for ITS-1/5.8S/ITS-2 rRNA ONT nemabiome metabarcoding when applied to fecal DNA

#### 3.3.1. Use of NC5 and NC2 primers for ITS-1/5.8S/ITS-2 rRNA ONT nemabiome metabarcoding directly on ovine fecal DNA leads to a large proportion of off-target reads

We had previously optimized the NC5/NC2 primers for ITS-1/5.8S/ITS-2 rRNA ONT metabarcoding on purified GIN L1 larvae with minimal contamination with fecal material. We next tested the performance on ovine fecal DNA. Genomic DNA was extracted from pooled fecal samples from nine flocks using repeated freeze–thaw cycles in liquid nitrogen followed by the *QIAamp PowerFecal Pro DNA Kit*. Amplification with NC5-Np-Adp and NC2-Np-Adp primers, followed by MinION Mk1B sequencing, generated 31,807 reads, of which only 47.4% mapped to nematodes (Supplementary Figure3). Off-target reads were a particular problem low-egg-count samples and were predominantly fungal (Supplementary Figure3B & C). Increasing annealing temperatures did not improve specificity (data not shown).

In silico analysis confirmed poor specificity: NC5 showed >80% identity to 8.2% of fungal 18S sequences (100% in 3.9%), while NC2 aligned to 37.4% of fungal 28S sequences (100% in 8.1%) (Supplementary Table 4). DNA prepared without the freeze–thaw step (using only the *QIAamp* kit) produced similar results but with slightly reduced nematode recovery (41.3% vs. 47.4% - data not shown).

#### 3.3.2 Development of new primers – EC1 and EC2 – to minimize off-target amplification when used for ONT ITS-1/5.8S/ITS-2 rRNA metabarcoding directly on ovine fecal DNA

Due to the substantial fungal co-amplification observed with NC5/NC2 primers when Nanopore nemabiome metabarcoding was directly applied to fecal stool DNA, we designed new primers, EC1 (forward) and EC2 (reverse), targeting conserved regions of nematode 18S and 28S rRNA while minimizing similarity to fungi (Figure 6; Supplementary Table 4).

**Figure 6.**
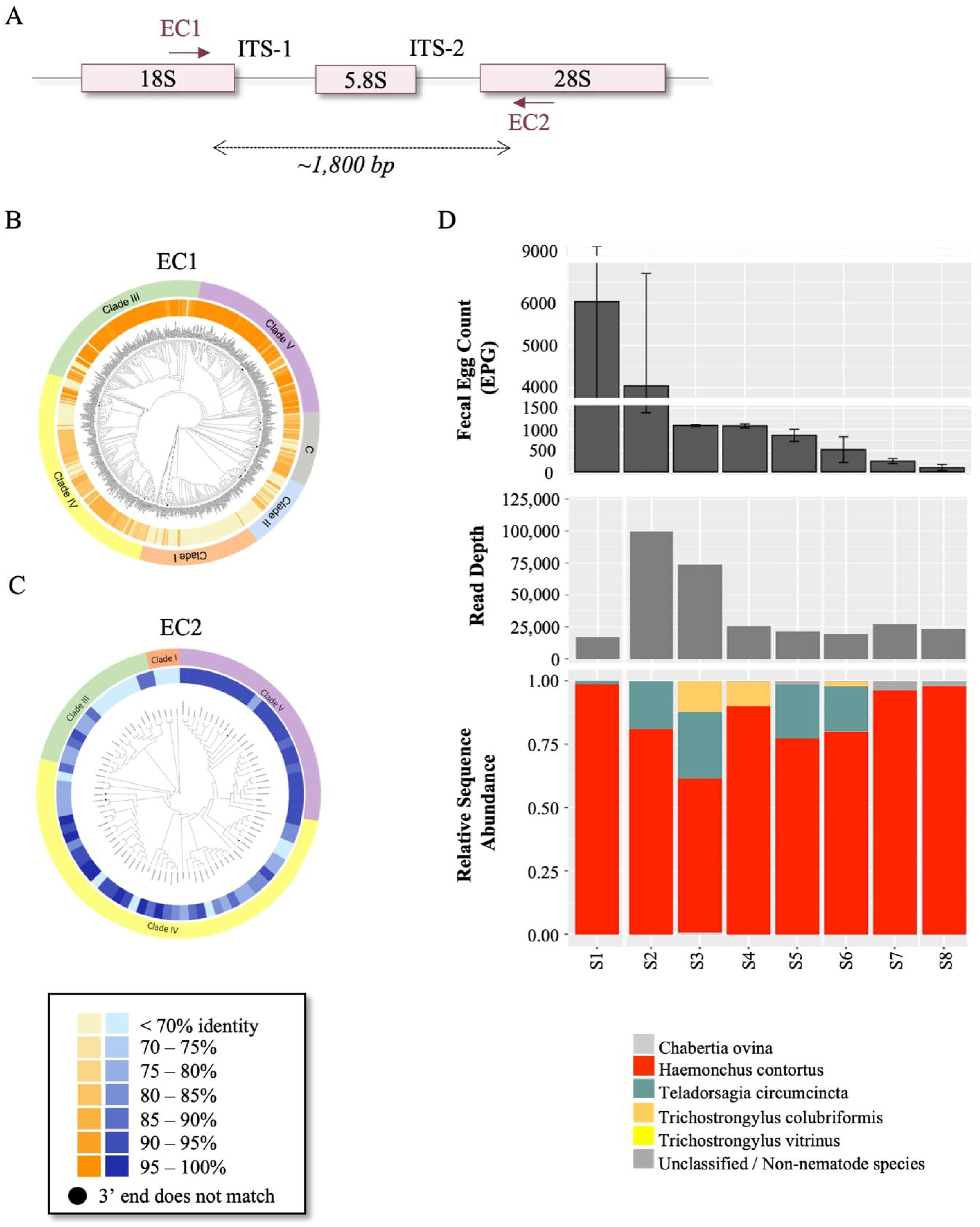
Design of the EC1/EC2 primer pair and assessment of their specificity for ITS-1/5.8S/ITS-2 metabarcoding when applied to fecal stool DNA. (A) Schematic representation of locus and primer positions. Using the PrimerTC tool and specific databases (18S or 28S / nemabiome.ca: https://staging--nemabiome.netlify.app/), we visualized the conservation of sequence identity of EC1 and EC2 primers with target reference sequence across the nematode phylum. In each diagram, the central phylogenetic tree represents a maximum likelihood tree of full-length sequences from the 18S rRNA database. Unique 18S trees were created for each diagram by exclusively incorporating the genera present in the databases used for PrimerTC analysis; (B) 18S, and (C) 28S. A black dot on the tree indicates a discrepancy between the last base pair of the primer sequence and the reference sequences of depicted genera in the database. The inner circle displays a heatmap, revealing the percentage identity between the tested primer and each genus in the chosen database, assessed with the PrimerTC tool. The outer circle signifies the Clade affiliation of nematode genera. (D) Primers EC1-Np-ADP and EC2-Np-ADP were used to amplify the ITS-1/5.8S/ITS-2 locus for Nanopore metabarcoding from genomic DNA prepared directly from fecal stool (without prior parasite extraction) using the “extended freeze/thaw protocol” and QIAmp Power Fecal Pro DNA Kit’ (materials and methods). The fecal samples were from nine different commercial sheep flocks with varying levels of GIN infection. The top histogram shows the fecal egg count of each sample (mean of 3 aliquots) with standard deviation. The middle histogram shows the number of mapped reads obtained for each sample. The lower bar chart shows the proportions of sequence reads mapping to different nematode species and non-nematode species (all non-nematode species reported as a single group represented in grey).

In silico analysis with PrimerTC showed that EC1 had >80% sequence identity with 70.8% of 2,645 nematode 18S reference sequences, with strong conservation in Clades III, IV, and V (Figure 6B). EC2 showed >80% identity in 83% of 254 nematode 28S sequences, with high conservation in Clades IV and V (Figure 6C). Together, the primers were predicted to provide broad coverage of Clade IV and V nematodes, which include the major ovine GIN species, but less complete coverage of other clades. Fungal cross-reactivity was minimal: EC1 and EC2 aligned to only 9.4% and 1.2% of fungal rDNA sequences, with no cases of 100% identity (Supplementary Table 4).

We validated EC1/EC2 experimentally using fecal DNA from eight sheep flocks (fecal egg counts ranging 100–6030 epg) which was extracted with the extended freeze–thaw plus *QIAamp PowerFecal Pro DNA Kit* protocol. Amplification with EC1-Np-Adp/EC2-Np-Adp primers and MinION Mk1B sequencing (10h run) produced 265,499 mapped reads, of which 91.8% were nematode sequences (Figure 6D). This suggests the EC1/EC2 primers will be more tolerant for metabarcoding applied to parasite samples with substantial fecal material contamination and also for the potential future application of ONT ITS-1/5.8S/ITS-2 rRNA metabarcoding directly to fecal DNA.

## Discussion

ITS-2 Illumina short-read nemabiome metabarcoding is well validated and increasingly used to characterize the species composition of gastrointestinal nematode communities (Avramenko et al. 2015; 2017; Redman et al. 2019; Queiroz et al. 2020; Beaumelle et al. 2021). Although a powerful technique, metabarcoding using paired-end Illumina short-read sequencing has several limitations. Firstly, it is restricted to markers of less than 600bp in length which can lead to limitations in discriminatory power and challenges with primer design. Secondly, the Illumina sequencing platforms are relatively inflexible; their fixed sequencing capacity means a high cost per unit unless a large number of samples are sequenced at once and sequencing runs take a relatively long time. Thirdly, the Illumina sequencing platforms require significant capital investment and aren’t often locally available in smaller laboratories. ONT sequencing potentially addresses these limitations as it allows long-read sequencing (up to 4.2 Mb) (https://nanoporetech.com/applications/investigations/genome-assembly), has a more flexible format for varied sample numbers and faster sequencing run times and requires little capital investment.

The goal of this study was to develop and validate ONT long-read metabarcoding to characterize GIN communities as a more flexible approach than Illumina short-read metabarcoding. We chose ovine GIN as they are a good system to develop and validate new metabarcoding methods for a number of reasons. GIN are a major cause of disease and production loss for the sheep industry worldwide and the rise of anthelmintic resistance means they are increasingly difficult to control (Chartier et al. 1998; Mortensen et al. 2003; Thomaz-Soccol et al. 2004; Cringoli et al. 2007; Sutherland et al. 2008; Burgess et al. 2012; Redman et al. 2015; Bosco et al. 2020; Melville et al. 2020). Consequently, GIN have been more intensively studied in sheep than in any other host. In addition, sheep GIN communities are predominantly composed of nematodes from the order Strongylida, most of which have reference sequences available for different parts of the rRNA cistron and curated databases are available for these at www.nemabiome.ca (O’Connor, Walkden-Brown, and Kahn 2006; Besier et al. 2016; Redman et al. 2019; Queiroz et al. 2020, Charrier et al, 2024; Workentine et al, 2020). The methods described here should also be relevant to ruminant livestock GIN more generally, including in cattle, goats and bison, as they have similar groups of GIN parasite communities.

### 4.1 ITS-1/5.8S/ITS-2 rRNA as a marker for long-read ONT metabarcoding of GIN communities from sheep and other ruminant domestic livestock

We chose the ITS-1/5.8S/ITS-2 rRNA as the marker with which to develop ONT long-read metabarcoding for several reasons. There is a relatively large number of reference sequences for this marker, and we previously developed a database of unique full-length nematode ITS-1/5.8S/ITS-2 rRNA reference sequences (available at www.nemabiome.ca) (Charrier et al, 2024). Ovine GIN predominantly comprises members of the superfamily Trichostrongyloidea for which multiple species are often too phylogenetically closely related for discrimination using an 18S rRNA marker which is commonly used for general nematode taxonomy. The non-transcribed ITS-2 rRNA has a greater evolutionary rate of divergence and so has been the standard marker for the identification of trichostrongylid nematode species for many years (Avramenko et al. 2015; 2017; Redman et al. 2019; Queiroz et al. 2020). Although ITS2 allows reliable species identification for most of the more common trichostrongylid nematodes of domestic ruminants, in some cases species discrimination can rely on a very small number of SNPs (Ramünke et al. 2018). This can lead to ambiguity, particularly as more reference sequences from different geographical regions are acquired over time. For example, an analysis of available ITS2 reference sequences in published 2015 suggested there were 3 fixed SNPs in ITS2 between *H. contortus* and *H. placei* but subsequent analysis that included additional reference sequences subsequently submitted to Genbank suggested this is not always the case (Ramünke et al.2018).

Pairwise analysis of currently available ITS-2 and ITS-1/5.8S/ITS-2 references sequences reveals a greater number of nucleotide differences between the major relevant trichostrongylid nematode species for the longer marker which should lead to more robust species discrimination (Figure 2 and Supplementary Figure 2). For example, there is range of 19-50 nucleotide differences between ITS-1/5.8S/ITS-2 reference sequences for *T. axei* and *T. vitrinus* compared to 0-26 differences for ITS-2. Another example is a range of 55-110 nucleotide differences between ITS-1/5.8S/ITS-2 reference sequences for the cattle parasites *C. oncophora* and *C. punctata* compared to just 2-23 for ITS2 (Figure 2B and Supplementary Figure 2B). One example of relevance to this current study is the lack of our ability to confidently discriminate between *C. curticei* and *C. fuelleborni* by short-read ITS2 metabarcoding due to there being as little as 2 nucleotide differences between the few available ITS2 reference sequences for these species as has been previously noted (Beaumelle et al. 2025). The distribution of *C. fuelleborni* is generally considered limited to Africa, and is mainly present in African buffalo, and so it seems the detection in this study by ITS2 metabarcoding is likely a misidentification of *C. curticei* (Figure 5) (Beaumelle et al. 2025). More detailed analysis with additional genetic markers will be necessary to confirm this.

Although the ITS-1/5.8S/ITS-2 marker should give more reliable discrimination of the more closely related trichostrongylid nematodes of domestic livestock than ITS2, the choice of genetic marker for metabarcoding also depends on the representation of the target species of interest in the respective databases. ITS2 still has better representation for the trichostrongylid nematode group overall and so maybe still a better choice, or at least a complementary choice, for some studies and applications at present. An example of this from this current study is the lack of an ITS-1/5.8S/ITS-2 reference sequences for *C. curticei* (or *C. fuelleborni*) which appears to have resulted in this species being misclassified as *C. oncophora* by ITS-1/5.8S/ITS-2 metabarcoding as discussed further below.

### 4.2 Comparison of ITS-2 and ITS-1/5.8S/ITS-2 nemabiome metabarcoding data

We used GIN L1 populations from two relatively large, previously published, studies conducted in two geographically distant countries; the UK and Canada (Redman et al. 2019; Queiroz et al. 2020). The species composition of these parasite populations had already been characterized by ITS-2 rRNA nemabiome metabarcoding allowing us to assess the accuracy of ITS-1/5.8S/ITS-2 rRNA long-read nemabiome metabarcoding undertaken on a subset of 34 and 43 samples from the Western Canadian and UK studies respectively. The genomic DNA used as a template was an aliquot of the same genomic DNA preparations used for the original ITS-2 nemabiome metabarcoding allowing a direct comparison of the data.

The ITS-1/5.8S/ITS-2 rRNA metabarcoding results showed a high degree of repeatability across three independent PCR replicates (Figures 3A and 4). Also, there was remarkably good overall agreement between the ITS-1/5.8S/ITS-2 rRNA and the original ITS-2 metabarcoding data. (Figure 5 and Supplementary Figure 1). Lin’s agreement coefficients were very high (0.92-0.98) for the five most abundant GIN species (*H. contortus, T circumcincta, T. colubriformis, T. vitrinus and O. venulosum*) and there was high agreement for the next highest species *C. ovina* (0.79-0.82). In addition, there were fewer reads assigned only to the *Trichostrongylu*s genus level for the ITS-1/5.8S/ITS-2 rRNA than the ITS-2 metabarcoding (1.64% Compared to 0% respectively). This is because the ITS-2 metabarcoding analysis pipeline assigns reads to the genus level if there is low confidence at the species level, contrary to the current long-read metabarcoding analysis pipeline which assigns to the closest related species level. There was lower agreement and some discrepancies for some of the lower abundance species (eg. *C. curticei/C. fuelleborni* and *T. axei*) which we believe is due to the lack of corresponding reference sequences in the ITS-1/5.8S/ITS-2 database. *Cooperia curticei* was present at 0.88 % overall with ITS-2 metabarcoding but was absent in the ITS-1/5.8S/ITS-2 data. Conversely, *C. oncophora* was identified in the ITS-1/5.8S/ITS-2 data which was unexpected since this is a cattle parasite but was not present in the ITS-2 data (Figure 5). These discrepancies can be accounted for by the lack of a *C. curticei* ITS-1/5.8S/ITS-2 sequence in the database and leading to ONT reads incorrectly mapping to the nearest reference sequence present (*C. oncophora*). Similarly, whilst the overall abundance of *T. axei* in the ITS-2 data was 1.8% it was only 0.44% in the ITS-1/5.8S/ITS-2 data (Figure 5). This could be due to the much larger number of ITS-2 reference sequences for this species (43) than ITS-1/5.8S/ITS-2 reference sequences (8) in the databases leading to less reliable mapping for the ONT long-read metabarcoding.

In summary, these minor discrepancies illustrate the critical importance of the completeness of reference sequence databases for reliable species assignment of metabarcoding sequence reads. Although the ITS-1/5.8S/ITS-2 marker should provide better discrimination of closely related parasite species due to the potentially higher number of variable sites, this is offset in the case of some species by the incompleteness of the reference database. At the time of writing, the ITS-2 database contains 12,436 unique sequences representing 1,721 distinct species compared to 10,107 unique sequences, representing 1,322 species for the ITS-1/5.8S/ITS-2 database. So overall, ITS-2 metabarcoding may still provide more comprehensive species assignments than ITS-1/5.8S/ITS-2 metabarcoding particularly for less common GIN species, ITS-1/5.8S/ITS-2 database. This emphasizes the importance of generating more ITS-1/5.8S/ITS-2 reference sequences in the future to support long read ONT metabarcoding. That having been said, it is noteworthy that, although the ITS-1/5.8S/ITS-2 database has a smaller representation of unique nematode species, it does include 171 Clade V nematode species which are absent from the ITS-2 database. This makes the point that for the most comprehensive and reliable coverage, a combination of taxonomic markers could be used.

### 4.3 The importance of filtering criteria for ITS-1/5.8S/ITS-2 rRNA ONT metabarcoding data

One of the major factors that has limited the adoption of ONT sequencing for metabarcoding studies has been the higher sequencing error rates of this chemistry compared to Illumina short-read sequencing (Delahaye and Nicolas 2021). However, there have been major improvements in this over the past two years. The release of the kit V14 chemistry (https://nanoporetech.com/q20plus-chemistry) vastly improved the sequencing read quality, and when paired with R10 flow cells and the Guppy super accuracy basecalling, it achieves the generation of sequence data with >99% raw read accuracy and this continues to improve. In our ITS-2/5.8S/ITS-2 metabarcoding experiments, on average, 27% of the ONT sequence reads were removed by our filtering steps, but the reads that were kept had over 99% accuracy as estimated from the phread Qscore (data not shown). As this average accuracy is higher than the 97% threshold commonly used for species assignment in Illumina ITS-2 nemabiome metabarcoding, these results were of sufficient quality to use the ONT reads for species-level classification. Moreover, we expect that with the application of the recently released Oxford Nanopore kit 14 chemistry, the percentage of sequencing read lost due to inadequate quality will lower, therefore further strengthening the use of this technique for nematode taxonomic classification.

However, it is important to be aware that these ONT error rates are still higher than for Illumina sequencing which could potentially lead to species mis-assignments, particularly when interested in rarer and very closely related species. Consequently, we used some additional manual filtering of the data in which we discarded any sample with less than a total of 1,000 reads or any species with less than 200 reads in the data overall. The total number of species identified by the ONT ITS-1/5.8S/ITS-2 rRNA metabarcoding before this step was 27 whereas it was 10 after this step (Table 1). The 17 species removed all had extremely low overall read numbers (1 to 317) and were species not expected to be present in ovine samples. Consequently, they are almost certainly artifactual.

### 4.4 A new ONT ITS-1/5.8S/ITS-2 rRNA nemabiome metabarcoding primer pair to minimize off-target sequence reads from fecal DNA

Current nemabiome metabarcoding studies using ITS-2 Illumina sequencing typically rely on parasite stages harvested from feces (eggs, L1s) or L3s from coprocultures (Avramenko et al., 2015; 2017; Redman et al., 2019; Queiroz et al., 2020). Whilst the particular parasite samples used in this study contained minimal fecal debris, parasite samples used for metabarcoding can contain variable amounts of fecal contamination. Further, in the longer term, the ability to apply nemabiome metabarcoding directly to fecal DNA would simplify protocols and make the method more suitable for routine diagnostic applications. Off-target amplification, predominantly from micro-organisms in feces, is one of the major limiting factors to achieve this. Consequently, we tested the performance of the NC5/NC2 primers, which are the most widely used in nematodes to amplify the ITS-1/5.8S/ITS-2 region, when applied to ovine fecal DNA since this primer pair was originally validated on adult worm DNA (Gasser et al., 1993; Newton et al., 1998). We found extensive fungal co-amplification with these primers which led to a major reduction in nematode read depth with the ONT ITS-1/5.8S/ITS-2 metabarcoding (Supplementary Figure 3). PrimerTC analysis confirmed strong sequence identity of NC5 and NC2 with fungal rDNA, consistent with these results. To address this issue, we used PrimerTC to develop new primers (EC1/EC2) targeting conserved nematode 18S and 28S regions while minimizing fungal similarity. These produced minimal off-target reads when used for ONT ITS-1/5.8S/ITS2 metabarcoding directly on ovine fecal DNA even at relatively low fecal egg counts (Figure 6). Hence this new primer pair should be more robust for use on sub-optimal samples and also have potential for direct fecal DNA metabarcoding. It should be noted however, that PrimerTC analysis revealed relatively poor conservation of these primer sites in Clades I and II, indicating that multiple primer sets would be required for more comprehensive nemabiome metabarcoding for applications where nematodes outside Clades IV and V are of interest.

## Glossary

BSA: Bovine Serum Albumin
CCC: Concordance Correlation Coefficient
CQ: Dr. Camila Queiroz
CM: Dr. Camila Meira
ddPCR: Droplet Digital PCR
EC: Dr. Eléonore Charrier
EPG: Eggs Per Gram
ER: Dr. Elizabeth Redman
FEC: Fecal Egg Count
gDNA: Genomic DNA
GIN: Gastrointestinal Nematode(s)
ITS: Internal Transcribed Spacer
JG: Dr. John Gilleard
L1, L3: First-stage and third-stage larvae
ONT: Oxford Nanopore Technologies
PCR: Polymerase Chain Reaction
RC: Rebecca Chen
SA: Dr. Sawsan Ammar

## Authors Contribution

EC co-conceptualized the study and undertook bioinformatic pipeline development, data analysis, and visualization and drafted the manuscript. ER and CQ provided the DNA samples and the ITS-2 metabarcoding results. EC developed and optimized an ITS-1/5.8S/ITS-2 metabarcoding protocol using the Oxford Nanopore Technologies MinION sequencing platform. EC and RC co-wrote the code for the bioinformatic pipeline. EC, ER, CM and SA were responsible for the processing of fecal samples and fecal egg count. EC developed and optimized the new EC1 and EC2 sequencing primers. EC performed, analyzed, and visualized the Nanopore sequencing experiments. ER performed the Illumina ITS-2 sequencing experiments. EC was responsible for the analysis and visualization of Illumina sequencing data with input from ER. EC wrote the draft manuscript with critical input and revision from JG. JG co-conceptualized the project and provided overall supervision.

## Competing Interest

The authors declare no competing interests.

## Funding Sources

This work was supported by Results Driven Agricultural Research (RDAR) grants 2022F059R and 2022F059RC. NSERC Discovery Grant, RGPIN-2021-02489 and NIH grant 5R01AI153088. We are also grateful to the Alberta Lamb Producers (ALP) and their members for the continued support of this work.

## Supplementary Material

**Supplementary Figure 1. Lin’s Concordance Correlation Coefficient analysis comparing the ITS-1/5.8S/ITS-2 and ITS-2 metabarcoding results for each major GIN species.**

The ITS-1/5.8S/ITS-2 rDNA was independently amplified from each L1 pool three separate times to produce three independent replicate libraries each sequenced on the Oxford Nanopore MinION for 30 hours. Species-specific Lin’s Concordance Correlation Coefficient analysis of the pairwise comparisons of the ITS-1/5.8S/ITS-2 nemabiome metabarcoding data between each independent replicate library and the ITS-2 metabarcoding data.

**Supplementary Figure 2. Inter and intra-species variation of ITS-2 sequence identity.**

All full-length ITS-2 sequences, currently available in the ITS-2 database (Workentine et al. 2019), were compiled for each species. (A) The number of sequences for each species is indicated next to the species name. Sequences were aligned with MUSCLE alignment (Edgar 2004) using the default parameters using Geneious version 10.1.3. The mean percent sequence between each species is listed. (B) The lowest and highest number of nucleotide differences between sequences for each species. The number of unique full-length sequences for each species in the database is indicated in parenthesis next to the species name. Sequences were aligned with MUSCLE (Edgar 2004) using the default parameters using Geneious version 10.1.3. An absolute pairwise character difference matrix was generated using the *haplotypes::distance()* R package.

**Supplementary Figure 3. Amplification of ITS-1/5.8S/ITS-2 and ONT long-read metabarcoding using the NC5/NC2 PCR amplification primers applied to genomic DNA prepared directly from ovine fecal DNA samples.**

Primers NC5-Np-ADP and NC2-Np-ADP were used to amplify the ITS-1/5.8S/ITS-2 locus for Nanopore metabarcoding from genomic DNA prepared directly from fecal stool (without prior parasite extraction) using the “extended freeze/thaw protocol” and QIAmp Power Fecal Pro DNA Kit’ (materials and methods). The fecal samples were from nine different commercial sheep flocks with varying levels of GIN infection. Panel A: Schematic representation of locus and primer positions. Panel B: The top histogram shows the fecal egg count of each sample (mean of 3 aliquots) with standard deviation. The middle histogram shows the number of mapped reads obtained for each sample. The lower bar chart shows the proportions of sequence reads mapping to different nematode species and non-nematode species (all non-nematode species reported as a single group represented in grey). Panel C: The relative proportional abundance of the sequence reads mapping to “non-nematode” reference sequences as determined by BLASTn analysis using the NCBI database (these are the sequence reads indicated in grey in panel B but broken down by group of species in this chart).

**Supplementary Table 1. Nucleotide sequences of primers used in this study.**

**Supplementary Table 2. Minimum and maximum length of nematode species of interest sequences found in the ITS-1/5.8S/ITS-2 database (Charrier et.al, 2024).**

**Supplementary Table 3. Nematode species taxonomically identified, and sequencing read numbers obtained through ITS-2 nemabiome metabarcoding, before and after applying manual curation criteria.**

The before-manual curation criteria analysis species and sequencing reads were obtained through the ITS-2 nemabiome metabarcoding bioinformatic pipeline. The after-manual curation criteria results were obtained after applying manual curation criteria where species with less than 200 reads were removed.

**Supplementary Table 4. Evaluation of PCR amplification primers sequence identity to fungal rDNA target sequences using the PrimerTC tool**

PrimerTC (Charrier et al, 2024) was used to assess the sequence identity of the NC2, NC5 (Gasser et al. 1993; Newton et al. 1998), EC1 and EC2 primers against the 18S (NC5 and EC1), 28S (NC2 and EC2) fungal reference sequence databases (https://ftp.ncbi.nlm.nih.gov/refseq/TargetedLoci/Fungi/).

## Notes

### Competing Interest Statement

The authors have declared no competing interest.

