## Supplementary figures and images for "The development and validation of long-read ITS-1/5.8S/ITS-2 nemabiome metabarcoding for ovine gastrointestinal nematodes using Oxford Nanopore Technologies (ONT) sequencing"

### Supplementary Figure 1

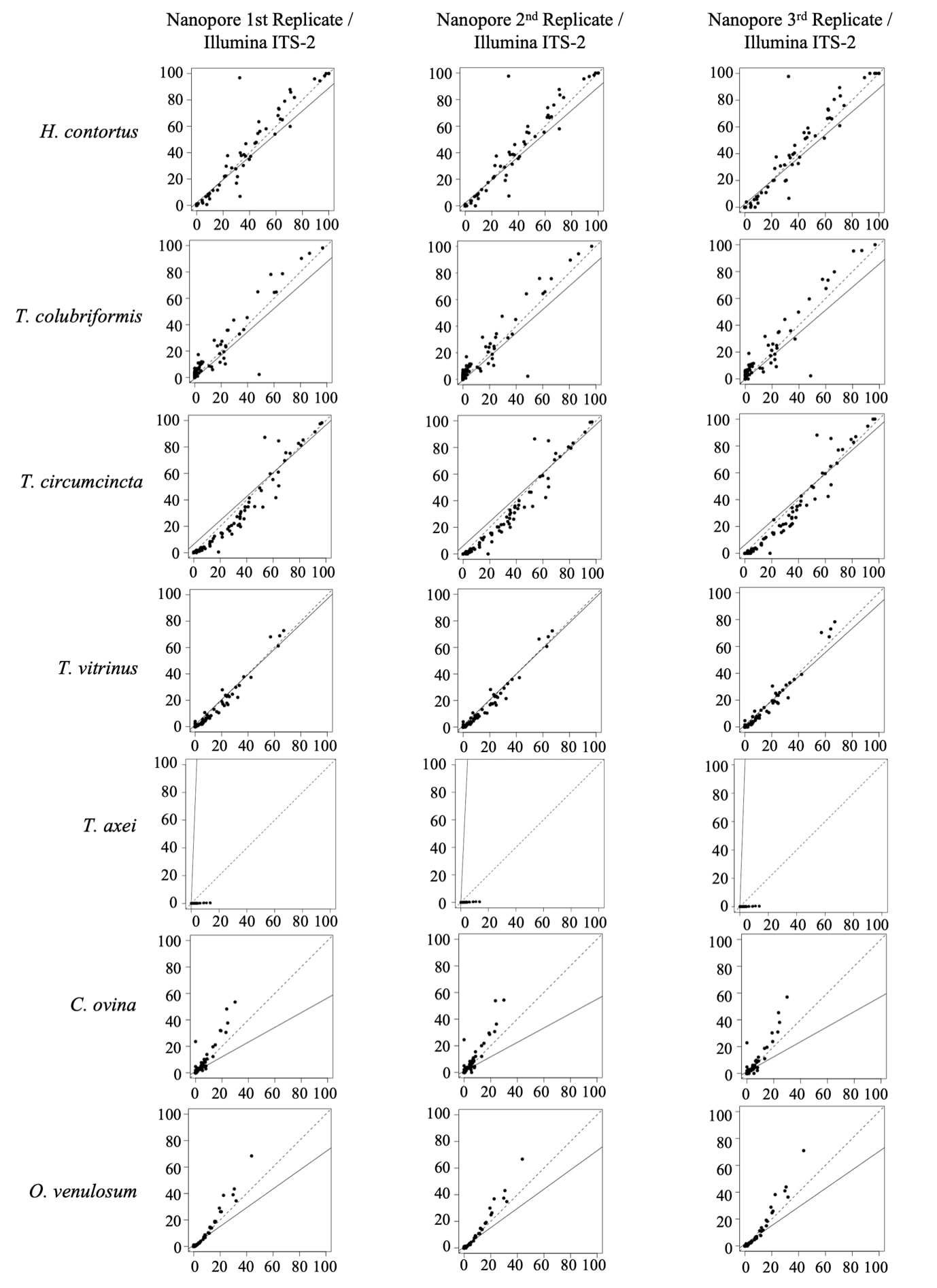

### Supplementary Figure 2

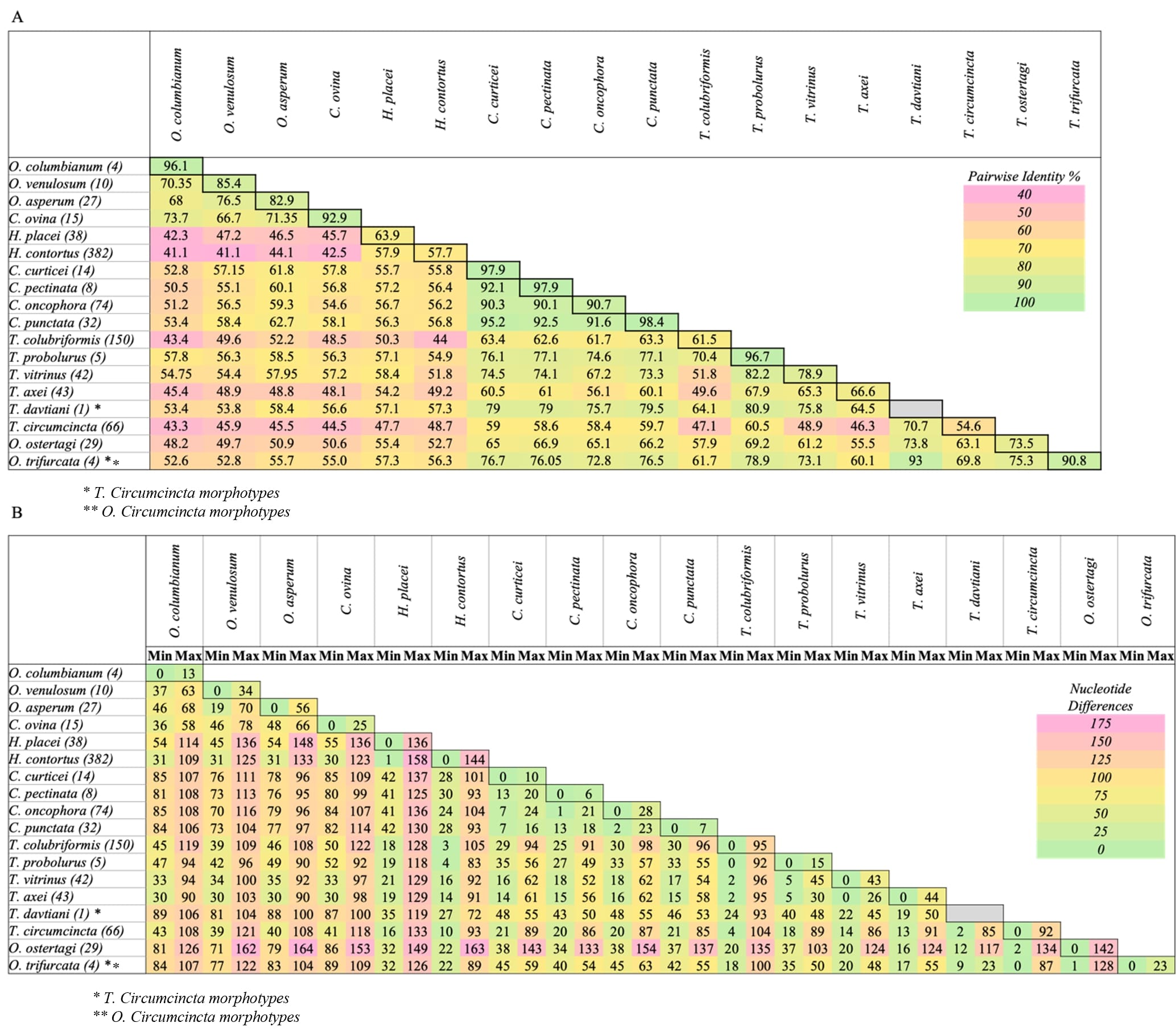

### Supplementary Figure 3

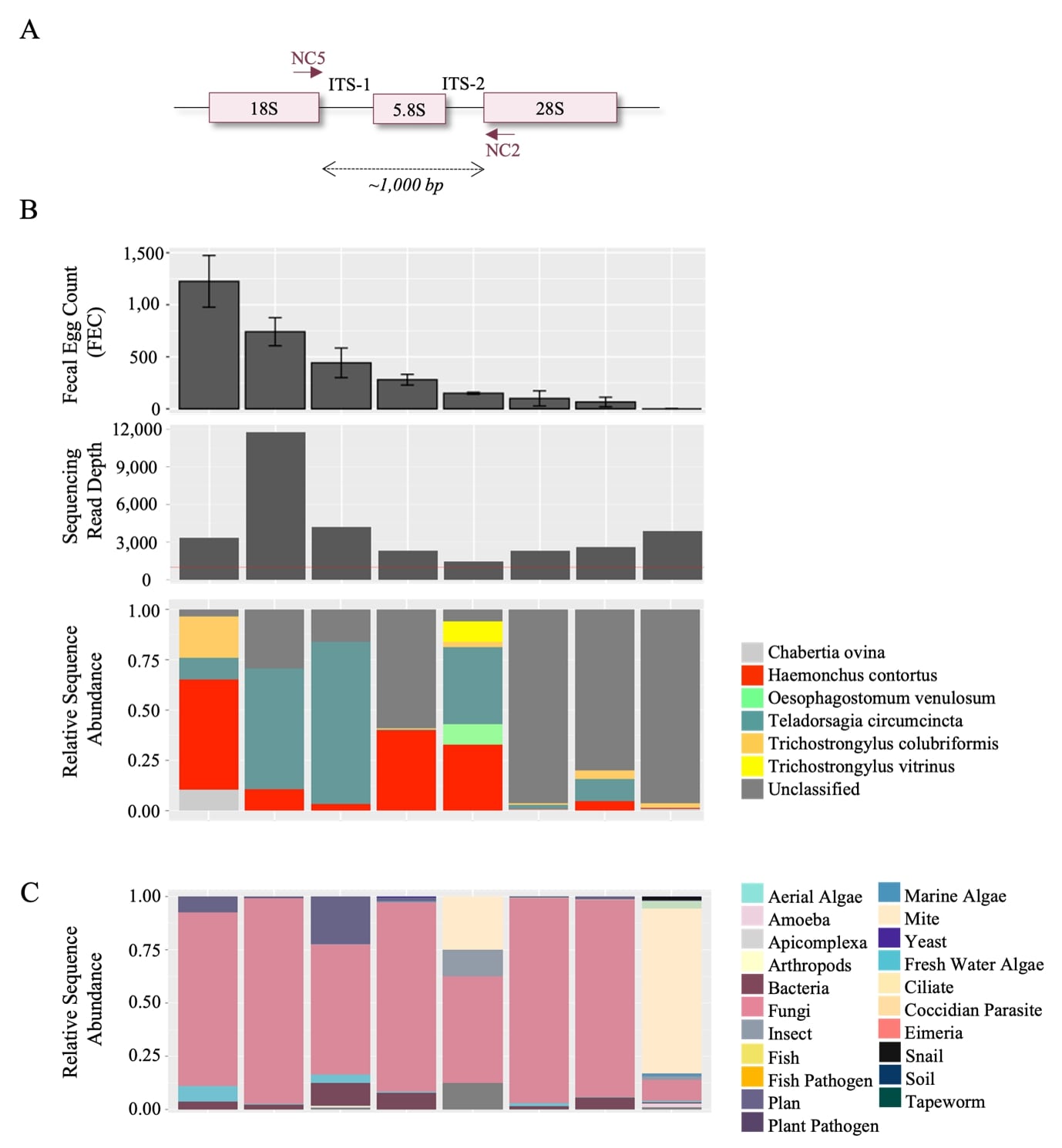

### Supplementary Table 1

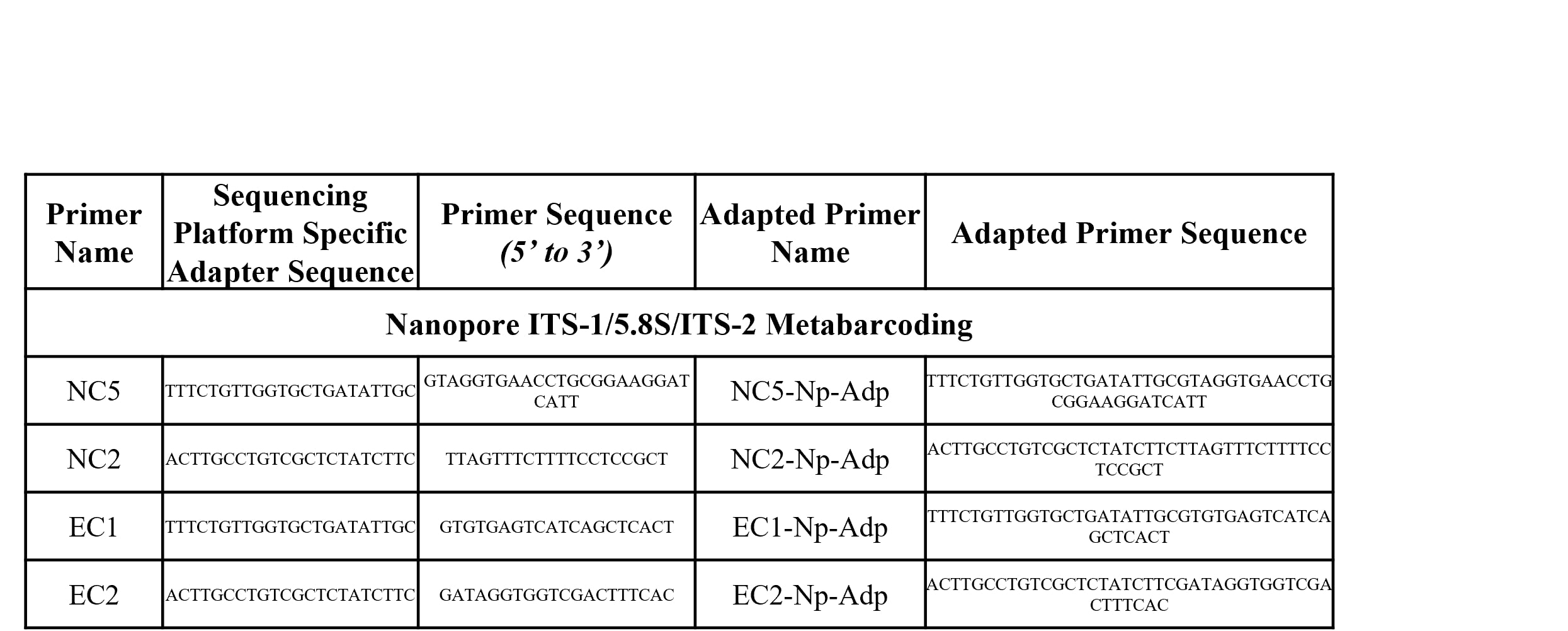

### Supplementary Table 2

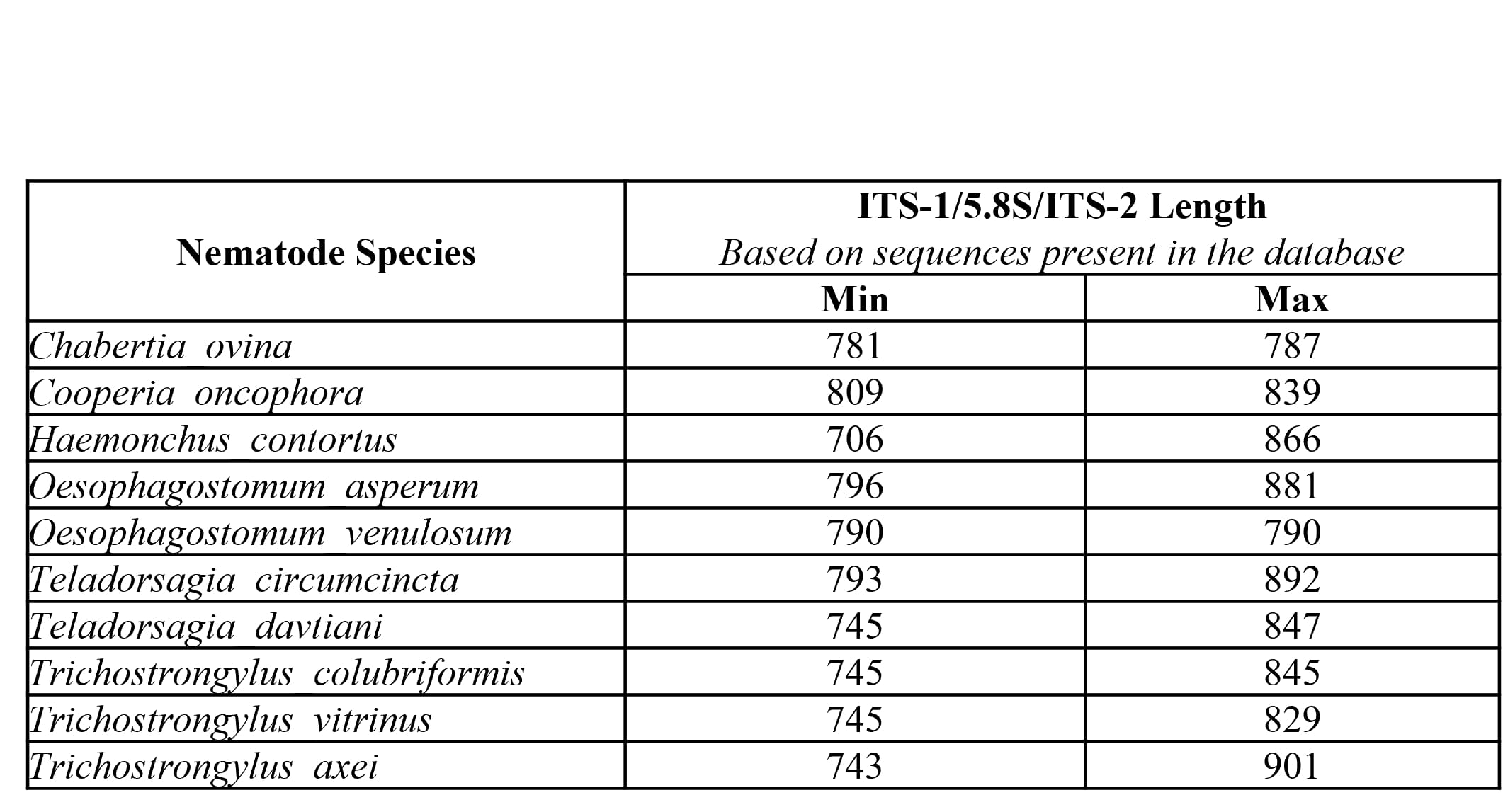

### Supplementary Table 3

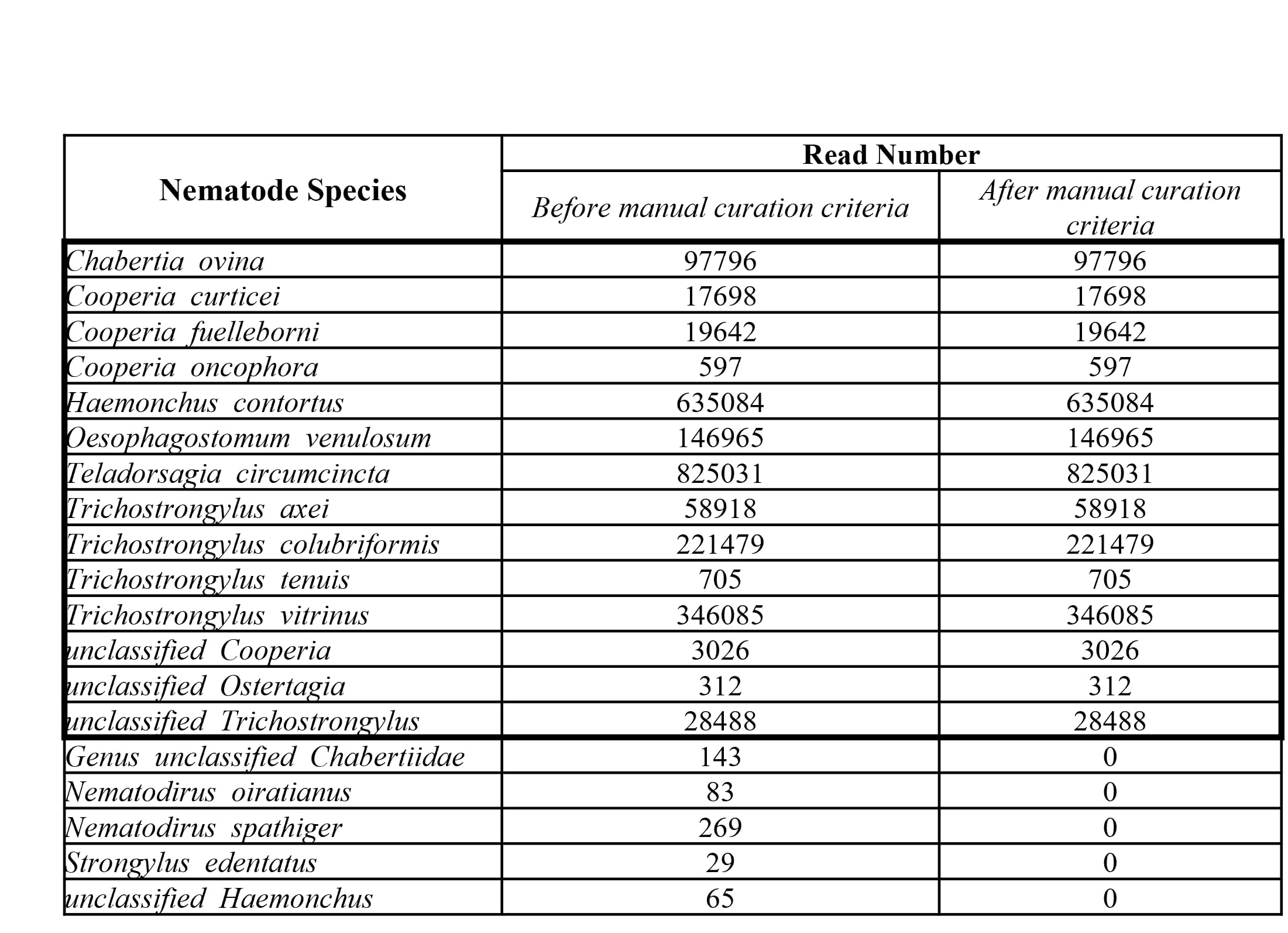

### Supplementary Table 4

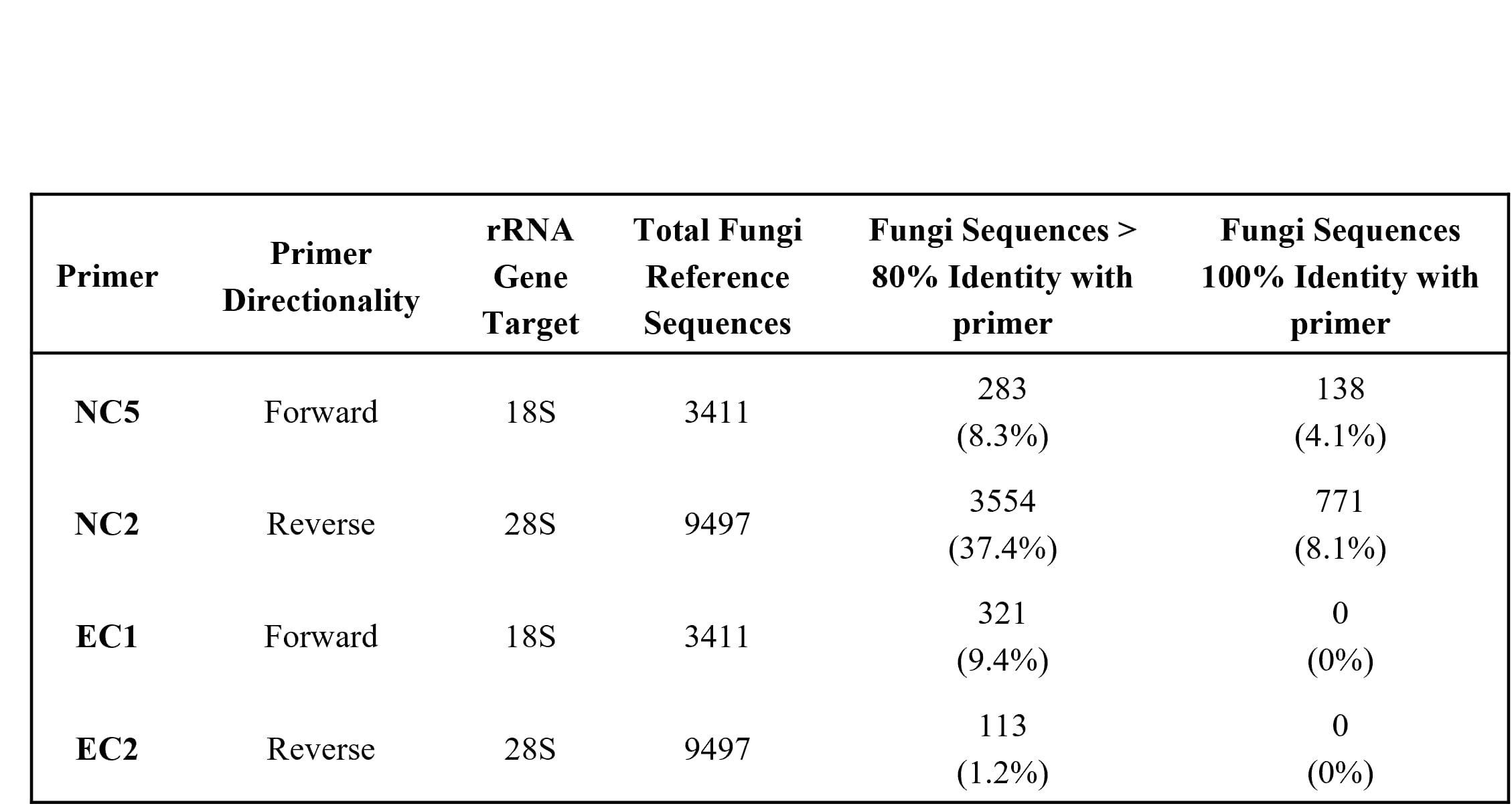
